# Conserved Chamber-Specific Polyploidy Maintains Heart Function in *Drosophila*

**DOI:** 10.1101/2023.02.10.528086

**Authors:** Archan Chakraborty, Nora G. Peterson, Juliet S. King, Ryan T. Gross, Michelle Mendiola Pla, Aatish Thennavan, Kevin C. Zhou, Sophia DeLuca, Nenad Bursac, Dawn E. Bowles, Matthew J. Wolf, Donald T. Fox

## Abstract

Developmentally programmed polyploidy (whole-genome-duplication) of cardiomyocytes is common across evolution. Functions of such polyploidy are essentially unknown. Here, we reveal roles for precise polyploidy levels in cardiac tissue. We highlight a conserved asymmetry in polyploidy level between cardiac chambers in *Drosophila* larvae and humans. In *Drosophila*, differential Insulin Receptor (InR) sensitivity leads the heart chamber to reach a higher ploidy/cell size relative to the aorta chamber. Cardiac ploidy-reduced animals exhibit reduced heart chamber size, stroke volume, cardiac output, and acceleration of circulating hemocytes. These *Drosophila* phenotypes mimic systemic human heart failure. Using human donor hearts, we reveal asymmetry in nuclear volume (ploidy) and insulin signaling between the left ventricle and atrium. Our results identify productive and likely conserved roles for polyploidy in cardiac chambers and suggest precise ploidy levels sculpt many developing tissues. These findings of productive cardiomyocyte polyploidy impact efforts to block developmental polyploidy to improve heart injury recovery.

## INTRODUCTION

Polyploidy (whole genome duplication) is a widespread property of somatic tissues and is present in at least 9 of 11 mammalian organ systems (Nandakumar et al., 2021; Peterson and Fox, 2021). Polyploidy is a common mechanism to generate larger cells (Marshall et al., 2012; Øvrebø and Edgar, 2018; Shu et al., 2018). While functions of tissue polyploidy largely remain mysterious (Fox et al., 2020), one potential clue is the local variation in tissue ploidy levels. For example, while all *Drosophila* midgut enterocytes and ovarian follicle cells are polyploid, the level of polyploidy varies along the A/P axis of these tissues (Lilly and Spradling, 1996; Viitanen et al., 2021). Similar examples of local polyploidy level variation are found in the fly nervous system, mouse liver, and mouse placenta (Nandakumar et al., 2020; Parisi et al., 2003; Tanami et al., 2017). Given the common relationship between ploidy level and cell size, controlled ploidy variation across a tissue could have profound impacts on tissue form and function.

Perhaps no animal cell type is more frequently polyploid than the cardiomyocyte. Developmental polyploidization of cardiomyocytes is highly conserved, occurring across mammals and avians, as well as in the fly *Drosophila* (Brodskiĭ, 1994; Derks and Bergmann, 2020; Hirose et al., 2019; Soonpaa et al., 1996; Yu et al., 2013). Such developmental increases in cardiomyocyte ploidy can involve increasing the number of nuclei per myocyte or by increasing the DNA content of mononucleate cardiomyocytes (Alkass et al., 2015; Bergmann et al., 2015; Brodskiĭ, 1994; Mollova et al., 2013; Peterson and Fox, 2021). Interestingly, developmentally programmed cardiomyocyte ploidy levels may vary by chamber, as suggested by comparisons of ventricular and atrial cardiomyocytes in mice and quail hearts (Anatskaya et al., 2001; Raulf et al., 2015). As summarized recently, the functional implications of developmentally acquired cardiomyocyte polyploidy are relatively unclear (Kirillova et al., 2021).

Moreover, a current focus of regenerative medicine research is to prevent developmentally programmed cardiomyocyte polyploidy. This focus comes from intriguing studies in mice and zebrafish that show that developmental polyploidization can block cardiac regeneration (González-Rosa et al., 2018; Han et al., 2020; Patterson et al., 2017). The interest in preventing developmental polyploidy is in line with findings that heart injury can cause cardiomyocytes to exceed polyploidy levels that are set by development and cause cardiac hypertrophy (Beltrami et al., 1997; Gan et al., 2020). However, the current focus on blocking developmental cardiomyocyte ploidy may prove problematic given the current lack of understanding of requirements for such naturally occurring whole genome duplication.

*Drosophila melanogaster* offers a highly tractable system to uncover conserved developmental regulation and functions of cardiomyocyte biology (Martínez-Morentin et al., 2015; Saha et al., 2022; Wessells et al., 2004; Wolf et al., 2006; Yu et al., 2013; Yu et al., 2015; Zechini et al., 2022). Flies and mammals share evolutionally conserved transcription factors that specify cardiac cell fate (Olson, 2006). Mutations in these genes cause heart defects in both humans and *Drosophila* (Akazawa and Komuro, 2005; Reim and Frasch, 2010; Stennard and Harvey, 2005). The cardiac organ in *Drosophila* is known as the dorsal vessel (Demerec, 1950; Rizki, 1978). In *Drosophila* embryos and larvae, this tubular organ consists of 104 cardiomyocytes spanning the A/P body axis from segments Thoracic 2 (T2) to Abdominal 7 (A7) (Molina and Cripps, 2001; Ocorr et al., 2007; Ponzielli et al., 2002; Zaffran et al., 1995, **Fig1A-A’)**. Based on embryonic expression of the Hox genes *Antennepedia (Antp), abdominal-A (abd-A), abdominal-B (abd-B)* and *Ultrabithorax* (*Ubx*), cardiomyocytes are specified into two distinct chambers along the anterior posterior axis: the aorta (T2-A4) and the heart (A5-A7). The aorta is further divided into the anterior aorta (T2-T3) and the posterior aorta (A1-A4) (Bataille et al., 2015; Lo and Frasch, 2001; Lovato et al., 2002; Monier et al., 2005; Perrin et al., 2004; Ponzielli et al., 2002; Schroeder et al., 2022). Each segment from A1 to A7 contains 8 cardiomyocytes separated by a pair of ostial cells, except for the posterior A7 segment (hereafter referred to as the apex) which consists of only 4 cardiomyocytes (Monier et al., 2005). A major function of the actively contracting larval cardiac organ is to disperse newly generated immune cells in the lymph (blood) throughout the animal (Johnstone and Cooper, 2006; LaBeau et al., 2009; Rotstein and Paululat, 2016; Rugendorff et al., 1994).This dispersal is accomplished by lymph perfusing into the heart chamber through ostia and then moving anteriorly through the heart and then aorta chambers (Babcock et al., 2008; Cevik et al., 2019; Curtis et al., 1999).

**Figure 1:**
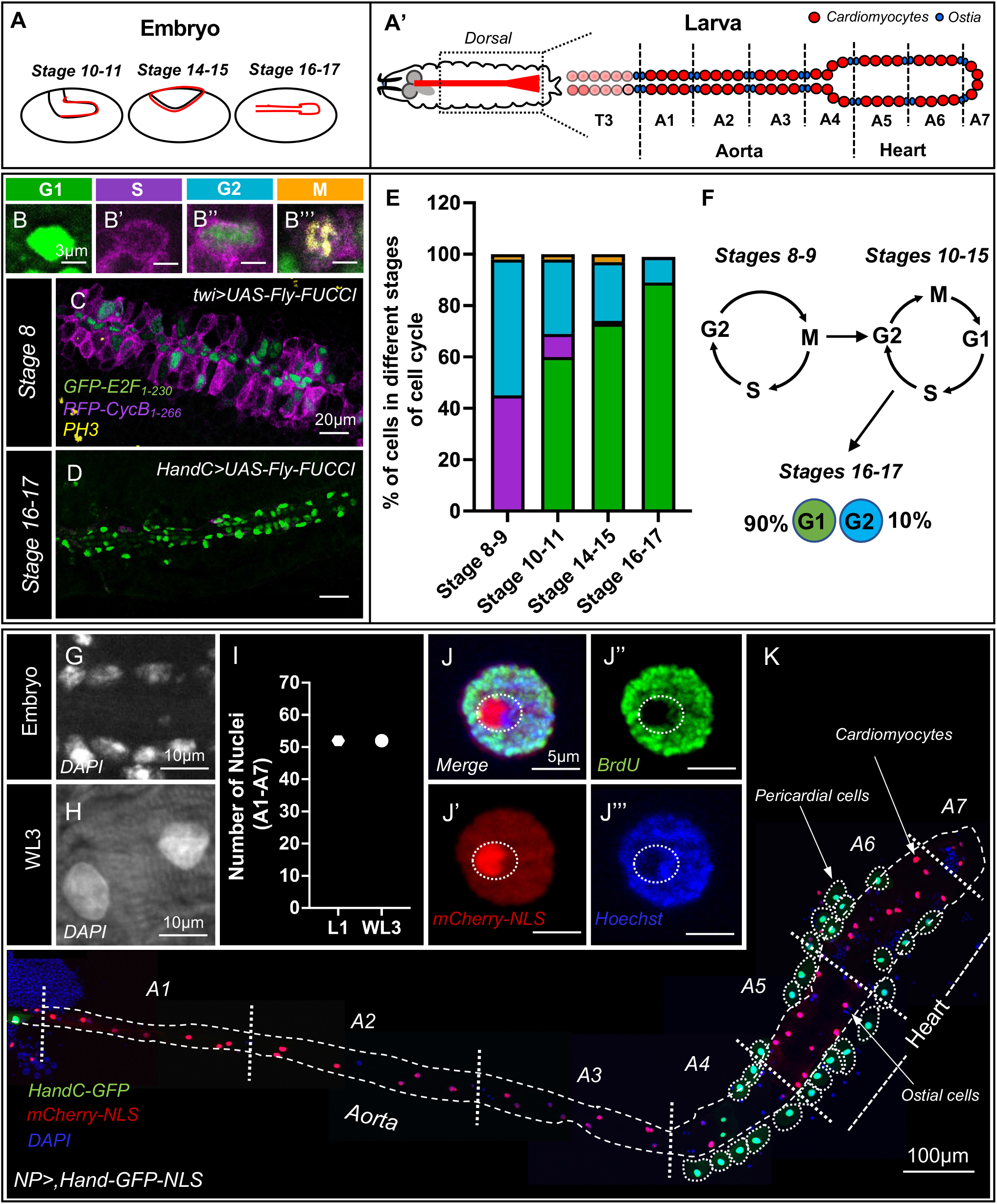
Embryonic and larval cell cycle activity of *Drosophila* cardiomyocytes. **(A-A’)** Schematics of the *Drosophila* cardiac organ (dorsal vessel, in red) during different stages of embryogenesis (**A**) and in larvae (**A’**). The dorsal vessel is divided into two distinct chambers along the anterior-posterior axis: the aorta and the heart. Segments A1 to A4 are referred to as the aorta and A5 to A6 are referred as heart. Segment A7 is referred to as the apex. From A1 to A7, each segment is separated by 2 pairs of ostial cells (blue) and consists of 8 cardiomyocytes (in red), except A7, which consists of only 4 cardiomyocytes. **(B-B’’’)** Examples of embryonic cardiomyocytes at different cell cycle stages labeled using Gal4-induced UAS-Fly-FUCCI. Scale bar = 20 μm. GFP-E2F1_1-230_ positive cells indicate G1 phase (**B**), RFP-CycB_1-266_ positive cells indicate S phase (**B’**), dual positive RFP-CycB_1-266_ and GFP-E2F1_1-230_ indicate G2 phase (**B’’**) and triple positive anti PH3, RFP-CycB_1-266_ and GFP-E2F1_1-230_ cells indicate M phase (**B’’’**). Scale bar = 3 μm. **(C)** Representative image of *twi-GAL4*+ stage 8 embryonic cardiomyocytes exhibiting active S and G2 phases. **(D)** Representative image of *HandC-Gal4+* stage 16-17 embryonic cardiomyocytes exhibiting G1 or G2 phase. **(E)** Quantitation of embryonic FUCCI data by stage. Drivers used for each stage: *twi-GAL4* for stage 8; *twi-GAL4* and *pnr-GAL4* for stage 9-11; *Mef2-GAL4* for stage14-15; and *Mef2-GAL4* and *HandC-GAL4* for stage 16-17. Colors for each cell cycle stage match the scheme in **B-B’’’**. n=10 embryos examined for each group. Each data set includes two or more biological repeats. (**F**) Schematic showing that embryonic cardiomyocytes undergo two types of mitotic cycles during stages 8-15 and arrest at G1 or G2 state at stage 16-17. **(G-H)** DAPI labels cardiomyocyte nuclei in embryonic (**G**) and Wandering Larval 3^rd^ instar (WL3, **H**) stages. Scale bar = 10 μm. **(I)** Number of total cardiomyocytes from segments A1-A7 in the first instar larval (L1) and WL3 stages. n=10 for each group. Each data set includes two or more biological repeats. Cardiomyocyte nuclei were identified as positive for *UAS-mCherry-NLS* driven by *NP5169-Gal4*. **(J-J’’’)** Representative image of BrdU positive cardiomyocyte in the heart chamber at WL3 for *NP5169-Gal4>UAS-mCherry-NLS*. Cardiomyocyte nuclei are labeled with mCherry (red), anti-BrdU (green) and Hoechst (Blue). Dotted outline highlights heterochromatin. Scale bar = 5 μm. **(K)** Immunofluorescence (IF) of WL3 cardiac organ (dorsal vessel). Cardiomyocytes are labeled with *NP5169-Gal4>UAS-mCherry-NLS* (NP>, red), pericardial cells with *HandC-GFP* (green, indicated by dotted outlines) and all nuclei with DAPI (blue). Scale bar = 100 μm.

Previously, adult *Drosophila* cardiomyocytes were found to be polyploid (Yu et al., 2013), but the developmental regulation and function of such polyploidy remained unknown. Here, we determine the developmental origin and regulation of this polyploidy and identify functional deficits in cardiac ploidy-reduced animals. We reveal that the transition from embryo to larvae coincides with cardiomyocytes entering G/S cycles known as endocycles (Schoenfelder and Fox, 2015; Shu et al., 2018; Zielke et al., 2013). This finding mirrors the transition to polyploid cardiomyocytes in many adolescent mammals (Alkass et al., 2015; Bergmann et al., 2015; Walsh et al., 2010). Further, we show that heart cardiomyocytes endocycle faster and reach a higher final ploidy and cell size than aorta cardiomyocytes. Differential Insulin Receptor (InR) signaling sensitivity underlies this chamber-specific asymmetry in cardiomyocyte ploidy. Reducing cardiomyocyte ploidy by knocking down *InR* or the endocycle regulator *fizzy-related* preferentially impacts the heart chamber compared to the aorta chamber. Such cardiac ploidy-reduced animals exhibit altered heart chamber dimensions, which are associated with diminished cardiac function, including stroke volume, cardiac output, and hemocyte velocity. As these defects in cardiac ploidy-reduced animals resemble systemic human heart failure, we examined chamber-specific ploidy and insulin signaling in human donor heart samples. These studies suggest that chamber-specific ploidy and insulin signaling levels are conserved between flies and humans. Overall, our studies highlight a role for precise, chamber-specific polyploidy in heart development and function, and caution against regenerative medicine strategies that block developmentally acquired cardiac polyploidy.

## RESULTS

### Cardiomyocytes undergo endocycles during larval development

To pinpoint the developmental onset of cardiomyocyte polyploidy in *Drosophila*, we first examined cardiac mesoderm cell cycle activity in the embryo (**Fig1A**). Many *Drosophila* cell types become polyploid in late embryogenesis by undergoing mitotic to endocycle transitions, while others initiate endocycles in early larval development (Edgar and Nijhout, 2004; Smith and Orr-Weaver, 1991). The entire mesoderm undergoes two divisions at embryonic stages 8 and 9, while more spatially distinct cardiac mesoderm divisions occur during late stage 10 and early stage 11, the time of cardiac fate commitment (Alvarez et al., 2003; Bate, 1993; Bodmer and Frasch, 1999; Foe, 1989; Han and Bodmer, 2003; Ward and Skeath, 2000). We used the mitotic marker Phospho-Histone H3 (PH3) along with *Gal4*-induced *UAS-Fly-FUCCI* (fluorescent ubiquitin-based cell cycle indicator, (Zielke et al., 2014a) to investigate cell cycle dynamics during these divisions (**Fig1B-E**).

As expression strength of commonly used cardiomyocyte lineage drivers differs at specific stages (**FigS1A**), we examined *UAS-Fly-FUCCI* driven by different drivers at different stages. Embryonic cardiac mesodermal cells are strongly *twist (twi)-GAL4+* (Greig and Akam, 1993) at stage 8, are *twi-GAL4+* and strongly *pannier* (*pnr)-GAL4+* (Heitzler et al., 1996) at stage 9 to 11, are strongly *Mef2-GAL4+* (Ranganayakulu et al., 1996) at stage 14-15, and are strongly *Mef2-GAL4+* and *HandC-GAL4+* (Albrecht et al., 2006) at stage 16-17 (**FigS1A**). Consistent with previous studies of embryonic cell cycle dynamics (Foe, 1989), stage 8 *twi-GAL4*+ mesodermal cells are either positive for RFP-CycB_1-266_ (S phase), dual positive for RFP-CycB_1-266_ and GFP-E2F1_1-230_ (G2 phase), or positive for PH3 (M phase). This expression pattern indicates S/G2/M cycles (**Fig1B-B’’’,C, E, F**). At stages 10-15, we observe these same cell cycle states in *pnr-Gal4*+ and *Mef2-GAL4*+ cardiac mesoderm, along with the addition of GFP-E2F1_1-230_ positive G1 phase. This expression pattern is consistent with a transition to G1/S/G2/M cycles (**Fig1B-B’’’, E, F**). Immediately after stage 15, both S and M phase states become undetectable in cardiac mesoderm. Instead, a majority (90%) of *HandC-GAL4*+ cells enter G1, while 10% enter G2 for the remainder of embryogenesis (**Fig1B-B’’’, D-F**). Given the co-occurrence of S phase and M phase (**Fig1E**) at every embryonic stage examined prior to stage 15, we conclude that embryonic cardiomyocytes do not endocycle, but instead undergo mitotic cycles followed by G1 or G2 arrest.

We next examined larval cardiomyocytes, organized into aorta and heart chambers (**Fig1A’**, Lovato et al., 2002; Molina and Cripps, 2001; Monier et al., 2005). By comparing cardiomyocyte nuclear size between late embryonic and late larval (Wandering Larval 3rd instar, WL3), we observe a clear increase (**Fig1G vs. H**). Over the same time period, we observe no increase in cardiomyocyte nuclear number (**Fig1I**). We find exactly 52 cardiomyocytes in both L1 and WL3 A1-7 segments, consistent with previous counts of late embryo cardiomyocyte nuclei (Ponzielli et al., 2002; Zaffran et al., 1995). These nuclear size and number comparisons support prior conclusions that cardiomyocytes are post-mitotic after embryogenesis (Molina and Cripps, 2001) and likely “polytenize”, a term associated with G/S cycles frequently termed endocycles (Stormo and Fox, 2017), during the larval stages (Rizki, 1978). We note that the lack of increased nuclear number from embryo to larva also suggests that cardiomyocytes do not undergo nuclear division followed by incomplete cytokinesis to become multinucleate.

To identify larval cardiomyocytes, we analyzed the expression patterns of several *Drosophila* cardiomyocyte drivers in both the heart and aorta chambers (**Fig1A’, FigS1**). Many commonly used cardiomyocyte drivers used for studies of the embryo or adult heart are weakly, regionally, or not expressed in the larval cardiac organ (**FigS1**). However, *NP5169-GAL4* (hereafter: *NP>*) and *Mef2-GAL4* (hereafter *Mef2>*, Molina and Cripps, 2001; Monier et al., 2005) are expressed in cardiomyocytes throughout the larval life cycle (**FigS1D-D’, G-G’**). As *Mef2*> also expresses in non-cardiac muscle cells (Nguyen et al., 1994; Taylor et al., 1995), we focused our study on *NP>*, which strongly and specifically expresses in all cardiomyocytes of segments A1-7 of the larval heart and posterior aorta (hereafter-aorta) but is not expressed in pericardial cells (**Fig1K**). In our analysis, we excluded thoracic cardiomyocytes (see Methods). To assess if cardiomyocytes re-enter the cell cycle during larval stages, we continuously fed larvae BrdU (see Methods) to identify S-phase activity from embryo hatching through all larval stages. From this feeding regimen, all WL3 cardiomyocytes are BrdU positive. All BrdU positive cardiomyocytes display an early S-phase pattern (**Fig1J-J’’’**), where heterochromatin (which is highly localized in the *Drosophila* nucleus, **Fig1J-J’’’**, area inside dotted outline) remains BrdU negative. This early-S only pattern is a hallmark of under-replication of heterochromatin that occurs in many endocycling *Drosophila* cells (Belyaeva et al., 1998; Lilly and Spradling, 1996; Sher et al., 2012; Smith and Orr-Weaver, 1991; Yarosh and Spradling, 2014). Overall, our results complete the picture of cell cycle dynamics in the embryonic and larval cardiomyocytes, revealing a progression from cell cycle arrest in late embryogenesis to larval endocycles.

### Polyploidy of larval cardiomyocytes is chamber-specific

We next examined the level of polyploidy in larval cardiomyocytes (segments A1-7). We used established microscopy methods to measure the ploidy (C) of *NP>+* cardiomyocytes in WL3 animals relative to a haploid 1C standard (see Methods). WL3 cardiomyocytes cluster into two populations with a median ploidy of ∼6C and ∼15C (**Fig2A**). The less than even genome doublings are consistent with our observation of under-replication (**Fig1J-J’’’**). Intriguingly, the two different populations of cardiomyocyte ploidy are mostly separated into distinct cardiac organ chambers (**Fig2A**). The higher ploidy population primarily corresponds to cardiomyocytes of the heart chamber, while the lower ploidy population primarily corresponds to the cardiomyocytes of the aorta chamber. Polyploidy frequently tracks with cell volume (Marshall et al., 2012; Schoenfelder and Fox, 2015). Notably, cell volume is reported to be higher in the heart chamber than the aorta, and the lumen of the heart is larger than that of the aorta (Lovato et al., 2002). In agreement, our sporadic FLP/FRT-mediated recombination (Xu and Rubin, 1993) labeling of mononucleated WL3 cardiomyocytes reveal a ∼1.6-fold larger area of heart vs. aorta cardiomyocytes (**Fig2B-D**). Taken together, we find that cardiomyocyte ploidy and cell size are higher in the heart chamber compared to the aorta (**Fig2E**).

**Figure 2:**
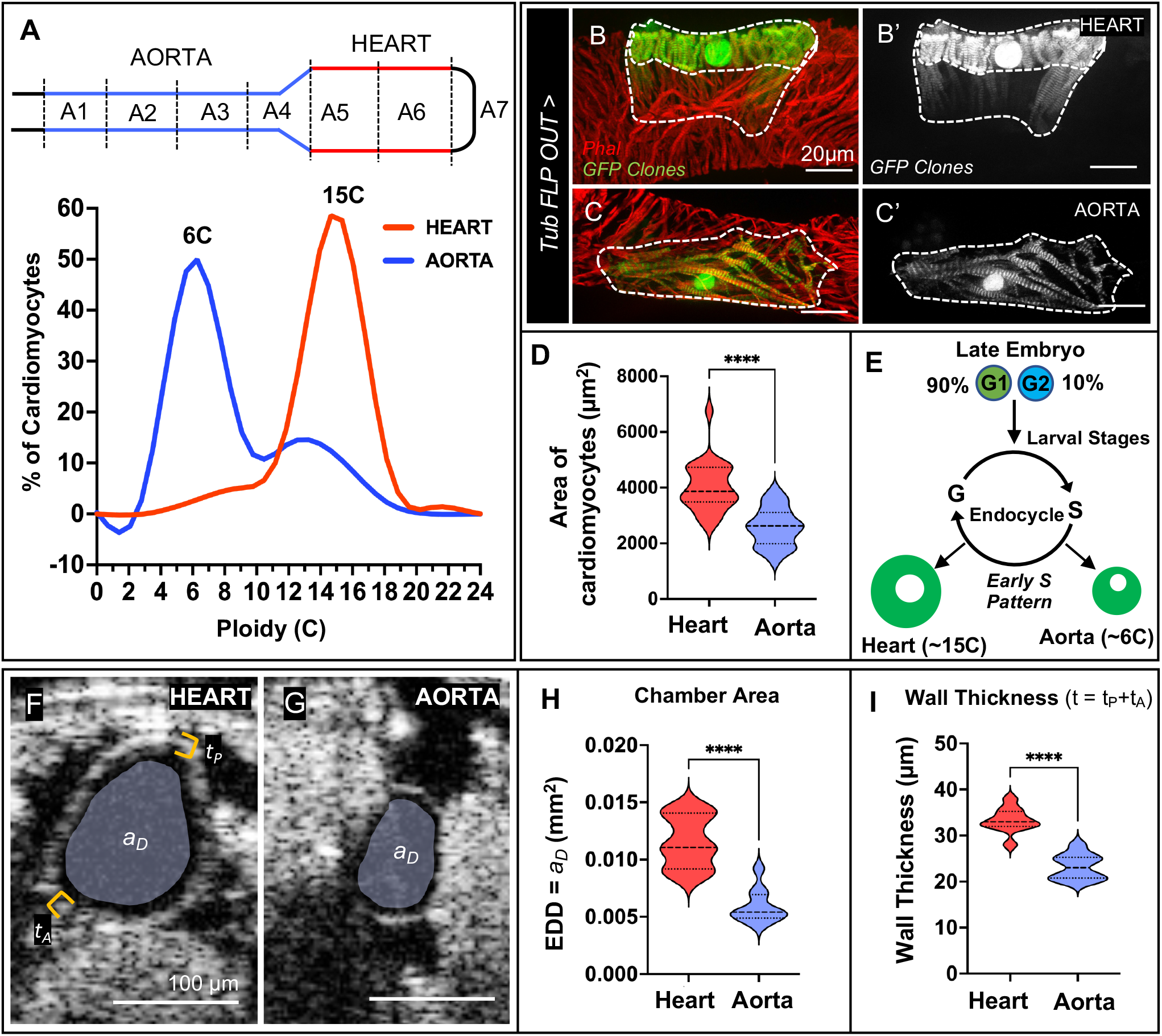
Polyploidization of *Drosophila* larval cardiomyocytes is chamber-specific. **(A)** Histogram showing ploidy distribution in the heart chamber (segments A5-A6, red line) and the aorta chamber (segments A1-A4, blue line) for *NP5169-Gal4>UAS-mCherry-NLS* (NP>). n=10 animals for each group. Each data set includes more than two biological repeats. **(B-C’)** GFP+ single cell FLP-out clones (green, see Methods) induced in the heart (**B-B’**) and aorta (**C-C’**) chambers. Dotted line highlights the periphery of the cardiomyocytes. Cardiac organ labeled with phalloidin (Phal, red). Scale bar = 20 μm. **(D)** Graph showing the single cell area of FLP-out GFP+ cardiomyocytes in the heart and aorta chambers. The results are shown as mean ± SD; ****P <0.0001 (Unpaired, two-tailed Student’s t-test). n=25 single cell clones for each group. Each data set includes more than two biological repeats. **(E)** Schematic showing that the larval cardiomyocyte endocycle program leads to chamber-specific ploidy levels. **(F-G)** Representative transverse two-dimensional real-time OCT images of the WL3 heart (**F**) and aorta (**G**) chamber End Diastolic Dimension area (EDD or a_D_, pseudo-colored in gray) for *NP5169-Gal4>UAS-mCherry* (NP>). Scale bar = 100 μm. Total wall thickness (t) is calculated as the sum of posterior (t_P_) and anterior (t_A_) wall thickness, indicated with yellow bracket. **(H)** Graph from OCT measurements of EDD (a_D_) in the heart and aorta chambers. The results are shown as mean ± SD; ****P <0.0001 (Unpaired, two-tailed Student’s t-test). n=10 for each group. Each data set includes two biological repeats. **(I)** Graph from OCT measurements of total wall thickness (t= t_A_+t_P_) in the heart and aorta chambers. The results are shown as mean ± SD; ****P <0.0001 (Unpaired, two-tailed Student’s t-test). n=10 for each group. Each data set includes two biological repeats.

We further analyzed ploidy within each chamber by cardiac organ segment at different temperatures and in different control genotypes. Overall chamber ploidy at 25°C is similar between two different control genotypes, namely *NP>* and *Mef2>* (**FigS2A**). Except for aorta segment A3, all aorta chamber segments (A1-A4) primarily have cells of the lower ploidy population (**FigS2B-E**). In contrast, cells of segments A5 and A6 of the heart chamber are almost exclusively of the higher ploidy population (**FigS2F, G**). We noticed minor but reproducible impacts on cardiomyocyte ploidy by temperature. Shifting animal culturing temperature from 25°C to 29°C slightly alters the distribution of ploidy (**FigS2A**). Upon closer examination, temperature slightly impacts ploidy in most cardiac organ segments except A3 and A4. (**FigS2B-H**). In A1, A2, and A7, ploidy slightly increases at 29°C, but in contrast in A5 and A6 ploidy slightly decreases at 29°C (**FigS2B-H**).

We next assessed if chamber-specific differences in cardiomyocyte ploidy and cell size are reflected in organ-level chamber size differences. We used transverse and sagittal optical coherence tomography (OCT) imaging (Choma et al., 2006; Wolf et al., 2006) in living WL3 animals. Compared to the lower ploidy aorta, the higher ploidy heart chamber exhibits a higher measure of lumen area known as the end diastolic dimension (EDD, *a*_*D*_, **Fig2F-H**) and a thicker chamber muscle wall (t, **Fig2I, FigS2I**, see Methods). Therefore, the higher ploidy heart chamber exhibits larger and thicker cardiomyocytes and an increased lumen area compared to that of the lower ploidy aorta. Our findings argue that endocycle regulation aids in sculpting chamber-specific differences in the larval cardiac organ.

### Heart cardiomyocytes endocycle faster than aorta cardiomyocytes due to differential insulin receptor regulation

To understand how endocycle regulation differs between the heart and aorta, we examined cardiomyocyte endocycle duration and timing. To do so, we performed a BrdU pulse-chase assay, examining S-phase activity over successive 24-hour time periods (see Methods). Immediately after hatching and continuing through the first 48 hours of larval development at 25°C, nearly every cardiomyocyte of the heart and the aorta actively undergoes DNA replication (**Fig3A-A’, B-B’, D**). However, this dramatically decreases at time points after 48 hours (**Fig3C-C’, D**). These results suggest that cardiomyocytes initiate endocycles at embryo hatching, during which ploidy increases. Interestingly, despite having a similar duration of endocycles (**Fig3D**), the heart chamber reaches a higher final ploidy than the aorta (**Fig2A**). Therefore, endocycles are faster in the heart relative to the aorta.

**Figure 3:**
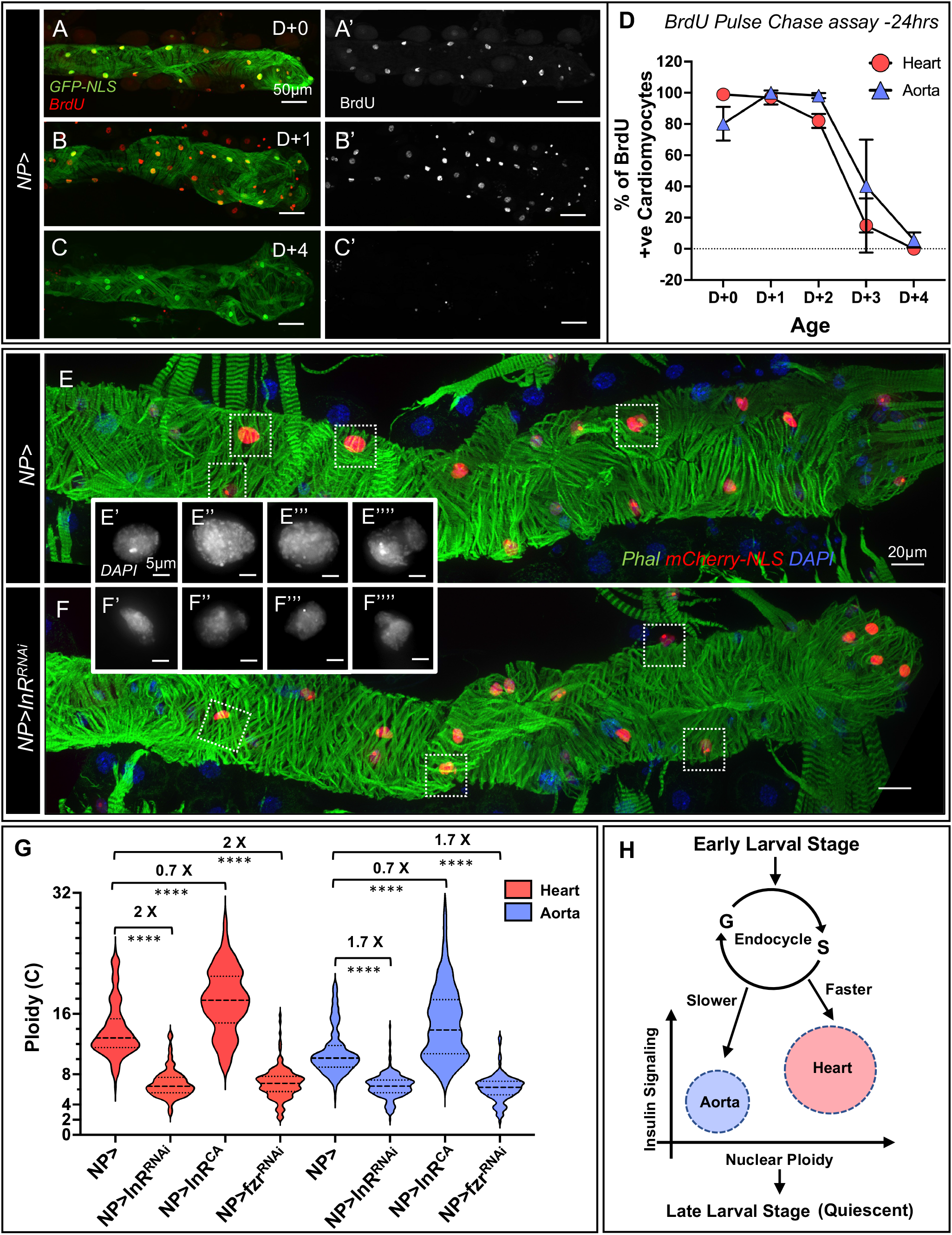
Cardiomyocytes in the heart chamber endocycle faster than in the aorta and are more sensitive to insulin signaling. **(A-C)** Representative images of BrdU positive (red) cardiomyocytes (green) in the WL3 heart chamber. *NP5169-Gal4>UAS-GFP-NLS* (NP>) animals were pulse-fed BrdU on control food (see Methods) for 24hrs at respective ages: D+0 (Embryos that hatched into larvae), D+1 (L1 larvae), D+2 (L2 larvae), D+3 (L3 larvae), D+4 (WL3 larvae). (**A-A’**) D+0, (**B-B’**) D+1, (**C-C’**) D+4. Scale bar = 50 μm **(D)** Graph showing the percentage of BrdU positive cardiomyocytes in the heart (red) and aorta (blue) chambers during larval development. n=5 animals for each group. Each data set includes two biological repeats. **(E-F)** Representative images of control *NP5169-Gal4, UAS-mCherry-NLS* X *w*^*1118*^ (NP>) and NP>*InR RNAi. UAS mCherry-NLS* (red) is expressed in all cardiomyocytes. The cardiac organ is also labeled with phalloidin (Phal, green) and DAPI (blue). Insets = enlarged view of single nuclei (from dotted boxes in lower magnification image) showing DAPI channel for *NP>* (**E’-E’’’’**) and *NP>InR RNAi* animals (**F’-F’’’’**). Scale bar = 20 μm; Inset scale bar = 5 μm **(G)** Graph of chamber-specific WL3 cardiomyocyte ploidy levels in each chamber of animals of the indicated genotypes, measured by microscopy (see Methods). A two-way ANOVA with multiple comparisons was used to compare chamber specific ploidy among each chamber in control (*NP>), NP>InR RNAi, NP>InR*^*CA*^ and *NP>fzr RNAi* animals and the results are shown as mean ± SD; ****P <0.0001. n= 10 for each group. Each data set includes two or more biological repeats. Numbers above each bracket indicate the fold change of the mean between groups. **(H)** Schematic illustration showing the timing of larval cardiomyocyte endocycles regulated by insulin signaling.

The early larval onset of cardiomyocyte polyploidy in *Drosophila* mirrors the early developmental transition to cardiomyocyte polyploidy in mammals (Bergmann et al., 2015). As in many larval *Drosophila* cells, this early larval ploidy increase coincides with the ability of the animal to take in food from the environment (Edgar and Nijhout, 2004; O’Farrell, 2004; Smith and Orr-Weaver, 1991). Indeed, starvation prevents cardiomyocyte endocycles (**FigS3A-D**). Diet influences PI3 Kinase/insulin signaling (Britton et al., 2002; Sudhakar et al., 2020; Zielke et al., 2011), which can regulate polyploidization, cell growth and cell proliferation (Bohni et al., 1999; Britton et al., 2002; Celton-Morizur et al., 2009; O’Brien et al., 2011; Saucedo et al., 2003; Shingleton et al., 2005; Shirakawa et al., 2017; Tamori and Deng, 2013; Zielke et al., 2011). Thus, we next assessed if InR activity regulates larval cardiomyocyte polyploidization. To do so, we used larval-onset *NP>* to drive *UAS-InR RNA interference (RNAi)*. Nuclear sizes of WL3 stage cardiomyocytes are visibly smaller in *NP>InR RNAi* animals compared to *NP>* controls (See Methods, **Fig3E, F**). Furthermore, cardiomyocyte DNA content decreases in *InR RNAi* animals (**Fig3E’-E’’’’, F’-F’’’’, G, FigS3E**). *NP>InR RNAi* does not impact cardiomyocyte nuclear number, indicating that our RNAi treatment does not impact embryonic mitotic cycles (**FigS3F**). Further, *NP>InR RNAi* does not alter *NP>* expression or noticeably change the pattern of actin-rich cardiac muscle striations, indicating that reduced InR signaling does not impact cardiac cell fate (**Fig3E vs. F**). Next, to test if InR activity is sufficient to drive endocycle activity, we constitutively expressed active InR (*NP>InR*^*CA*^) in larval cardiomyocytes. This treatment increases the overall ploidy of cardiomyocytes without altering cardiomyocyte number (**Fig3G, FigS3E-F**). These results show that InR activity is necessary and sufficient for WL3 cardiomyocytes to achieve final ploidy through endocycles.

To perturb polyploidy independently of insulin signaling, we additionally assessed if the well-known endocycle regulator and activator of the Anaphase Promoting Complex (APC), Fizzy-related (*fzr*)/Cdh1 (Sigrist and Lehner, 1997; Zielke et al., 2013), also controls cardiomyocyte ploidy. Indeed, *NP>fzr RNAi* cardiomyocytes exhibit significantly reduced ploidy (**Fig3G, FigS3E**). Importantly, *NP>fzr RNAi* does not impact cardiomyocyte nuclear number (**FigS3F**), confirming that our experimental conditions do not impact the known role for *fzr* in controlling embryonic cardiomyocyte cell division (Drechsler et al., 2018).

Although *NP>* is expressed well in both the aorta and heart (**Fig1K**), our genetic manipulations using this driver revealed more profound impacts on ploidy in the heart chamber. The mean ploidy of the A5 and A6 heart chamber segments is reduced 2.0-fold in *NP>InR RNAi* and *NP>fzr RNAi* WL3 animals, while ploidy in A1-4 aorta segments is reduced only 1.7-fold (**Fig3G, FigS3H, I, K**). These results suggest that the heart chamber may upregulate endocycle machinery to a greater extent than the aorta to achieve faster endocycles and a higher final ploidy. In agreement, experimentally elevating InR in both chambers (*NP>InR*^*CA*^) increases median ploidy equally (by 0.7-fold from controls in each chamber, **Fig3G, FigS3J**). Overall, these results highlight differences in InR sensitivity in cardiomyocytes that underlie ploidy differences between two cardiac organ chambers (**Fig3H**). Of the two chambers, the heart is the most impacted by loss of InR and achieves the higher final ploidy.

### Heart chamber size is particularly sensitive to ploidy reduction

We next examined cardiac organ dimensions in animals with altered cardiac ploidy. For ploidy reduction, we used both *InR* and *fzr RNAi* animals, as InR can also have ploidy-independent roles (Saltiel and Kahn, 2001; Yoon, 2017), Notably, our manipulations of *fzr* and *InR* cell cycle and growth signals do not alter body weight (**FigS3G**), suggesting that we are not substantially altering overall body dimensions. Using OCT in live WL3 animals, we next investigated chamber-specific dimensions of the cardiac organ with reduced (*NP>InR RNAi* and *NP>fzr RNAi*) or increased (*NP>InR*^*CA*^) ploidy. We measured chamber length (*l*), wall thickness (t), the end-diastolic dimension (EDD, *a*_*D*_) when the heart is relaxed, and the end-systolic dimension (ESD, *a*_*S*_) when the heart is contracted (**FigS2I**, see Methods). Previously, changes in wall thickness in a model of *Raf*-induced cardiac hypertrophy were shown to be independent of ploidy level (Yu et al., 2013; Yu et al., 2015). Similarly, our larval ploidy manipulations (both increasing and decreasing cardiomyocyte ploidy) do not impact wall thickness in either the heart or aorta (**FigS4A**).

Although our experimental approach to alter cardiomyocyte ploidy had no impact on wall thickness, we found several size alterations in the heart chamber in cardiomyocyte ploidy-reduced animals. In both *NP>InR RNAi* and *NP>fzr RNAi* animals, the heart chamber length (**Fig4A, B, D- lH, Fig4E, Movie S1**) and end diastolic dimension area (EDD, a_D_, **Fig4F, G, I, J, Movie S1**) are significantly reduced. In contrast, the end systolic dimension area (ESD, a_S_) of the heart chamber is not affected in cardiac ploidy-reduced *NP>InR RNAi* and *NP>fzr RNAi* animals (**FigS4C-G**). From our length and area measurements, we computed stroke volume, which approximates the amount of hemolymph pumped during each systolic contraction (**FigS2I**, see Methods). Stroke volume of the heart chamber is significantly decreased for both *NP>InR RNAi* and *NP>fzr RNAi* animals (**Fig4P**). In contrast to the heart chamber, the aorta of *NP>InR RNAi* and *NP>fzr RNAi* animals remains similar to controls in terms of the same functional measurements. Specifically, we do not observe any significant change in a_D_, chamber length, and stroke volume of the aorta, except for a slightly reduced chamber length in *NP>fzr RNAi* animals (**Fig4K-O, FigS4B, M**). In cardiac ploidy-increased *NP>InR*^*CA*^ animals, there is no change in a_D_, a_S_, length, or stroke volume of the heart or aorta chambers (**Fig4C, E, H, J, M, O, P, FigS4B, E, G, J, L, M**). Therefore, our *NP>*-driven approach to increase larval cardiomyocyte ploidy by *InR*^*CA*^ did not cause noticeable cardiac hypertrophy. Overall, our results indicate that the heart, but not the aorta, of cardiac ploidy-reduced mutants exhibits numerous alterations in heart chamber size, which impact cardiac physiology. The preferential impact on the heart is consistent with this chamber’s higher ploidy and dependency on InR signaling.

**Figure 4:**
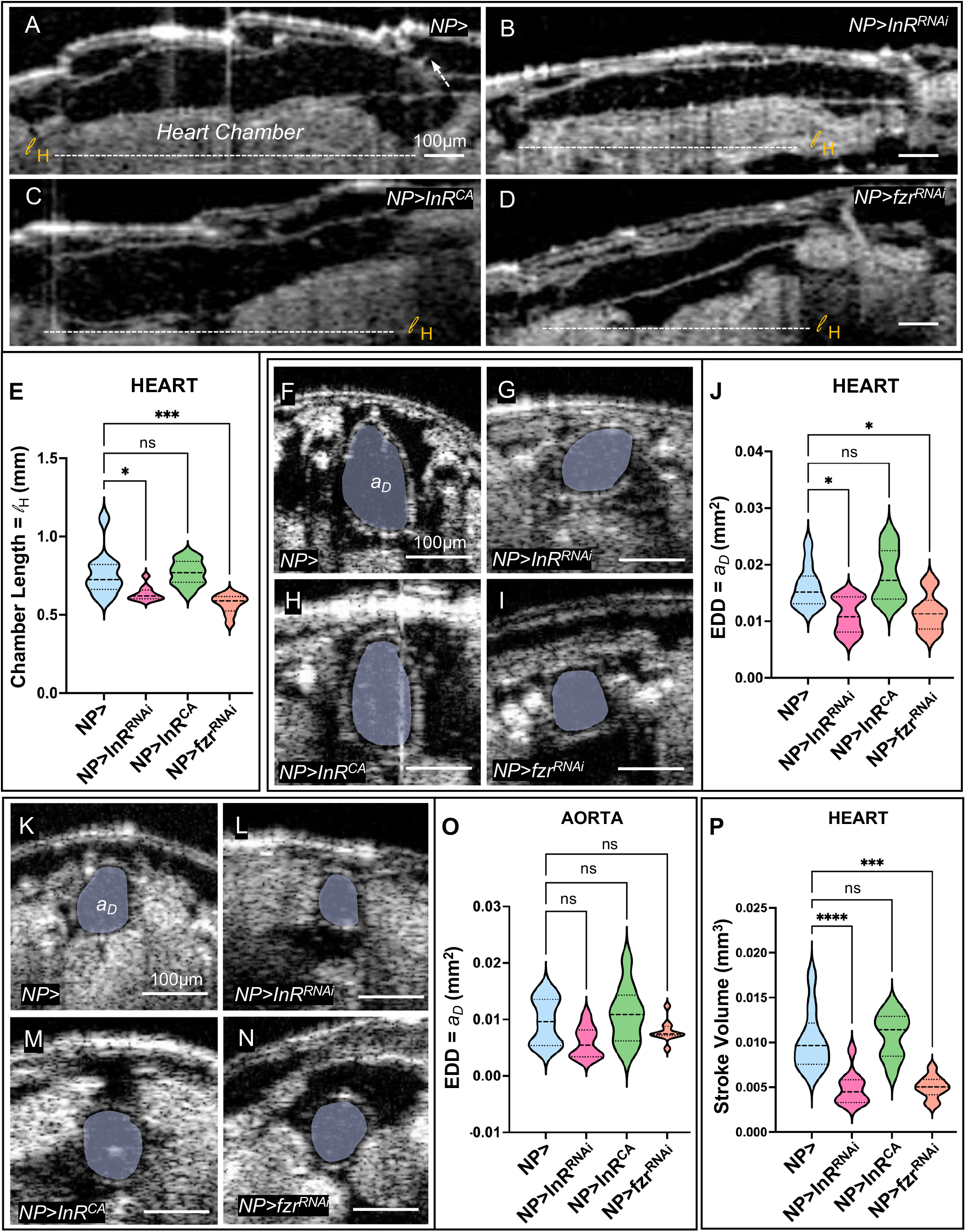
Size of the heart chamber is reduced in cardiac ploidy-reduced animals. **(A-D)** Representative sagittal two-dimensional real-time OCT images of the WL3 heart chamber of control *NP5169-Gal4, UAS-mCherry-NLS* X *w*^*1118*^ (*NP>)* (**A**), *NP>InR RNAi* (**B**), *NP>InR*^*CA*^ (**C**), and *NP>fzr RNAi* (**D**). White arrow in **A** indicates the posterior boundary of the heart chamber. White dashed line indicates the chamber length (l_H_). Scale bar = 100 μm **(E)** Graph showing heart chamber length in animals of the indicated genotype. A two-way ANOVA with multiple comparisons was used to compare OCT measurements of heart chamber length for control (*NP>), NP>InR RNAi, NP>InR*^*CA*^ and *NP>fzr RNAi* animals and the results are shown as mean ± SD; *P<0.05, ***P<0.001. n= 10 for each group. Each data set includes two biological repeats. **(F-I)** Representative transverse two-dimensional real-time OCT images of WL3 heart chamber End Diastolic Dimension area (EDD or a_D_, pseudo-colored in gray) for control *NP5169-Gal4, UAS-mCherry-NLS* X *w*^*1118*^ (*NP>)* (**F**), *NP>InR RNAi* (**G**), *NP>InR*^*CA*^ (**H**), and *NP>fzr RNAi* (**I**). Scale bar = 100 μm **(J)** Graph of a_D_ in the WL3 heart chamber in animals of the indicated genotypes. A two-way ANOVA with multiple comparisons was used to compare control (*NP>), NP>InR RNAi, NP>InR*^*CA*^ and *NP>fzr RNAi* animals and the results are shown as mean ± SD; *P<0.05. n= 10 for each group. Each data set includes two biological repeats. **(K-N)** Representative transverse two-dimensional real-time OCT images of a_D_ as in (**F-I**) but for the aorta chamber of control (*NP>)* (**K**), *NP>InR RNAi* (**L**), *NP>InR*^*CA*^ (**M**) and *NP>fzr RNAi* (**N**). Scale bar = 100 μm **(O)** Graph of aorta chamber a_D_ for animals of the indicated genotypes. A two-way ANOVA with multiple comparisons was used to compare OCT measurements of aorta chamber a_D_ for control (*NP>), NP>InR RNAi, NP>InR*^*CA*^ and *NP>fzr RNAi* animals and the results are shown as mean ± SD; ^ns^P>0.05. n= 10 for each group. Each data set includes two biological repeats. **(P)** Graph of stroke volume in the WL3 heart chamber of animals of the indicated genotypes. A two-way ANOVA with multiple comparisons was used to compare OCT measurements of heart chamber stroke volume for control *NP5169-Gal4, UAS-mCherry-NLS* X *w*^*1118*^ (*NP>), NP>InR RNAi, NP>InR*^*CA*^ and *NP>fzr RNAi* animals and the results are shown as mean ± SD; ***P<0.001, ****P<0.0001. n= 10 for each group. Each data set includes two biological repeats.

### Cardiac ploidy-reduced mutants have disrupted cardiac output and hemocyte movement

As the ultimate function of a cardiac organ is to regulate blood or lymph flow, we next measured cardiac output and hemocyte velocity in control and cardiac ploidy-reduced animals. Using OCT, we first determined the heart rate of *NP>InR RNAi* and *NP>fzr RNAi* WL3 animals. We measured the heart beats per minute for animals of each genotype. Heart rate is not significantly different in either chamber between control, *NP>InR RNAi* and *NP>fzr RNAi* WL3 animals (**Fig5A-D, FigS4N**). These results differ from findings of heart rate regulation by *InR* in adult *Drosophila* during aging (Wessells et al., 2004). We next calculated cardiac output (**FigS2I**), which is the product of stroke volume (**Fig4P, FigS4M**) and heart rate (**Fig5D, FigS4N**). Cardiac output of the heart chamber is significantly reduced in cardiac ploidy-reduced *NP>InR RNAi* and *NP>fzr RNAi* animals, but not in the aorta chamber (**Fig5E, F**). Our cardiac output results are consistent with cardiac ploidy-reduced animals exhibiting a compromised measure of heart (not aorta) function.

**Figure 5:**
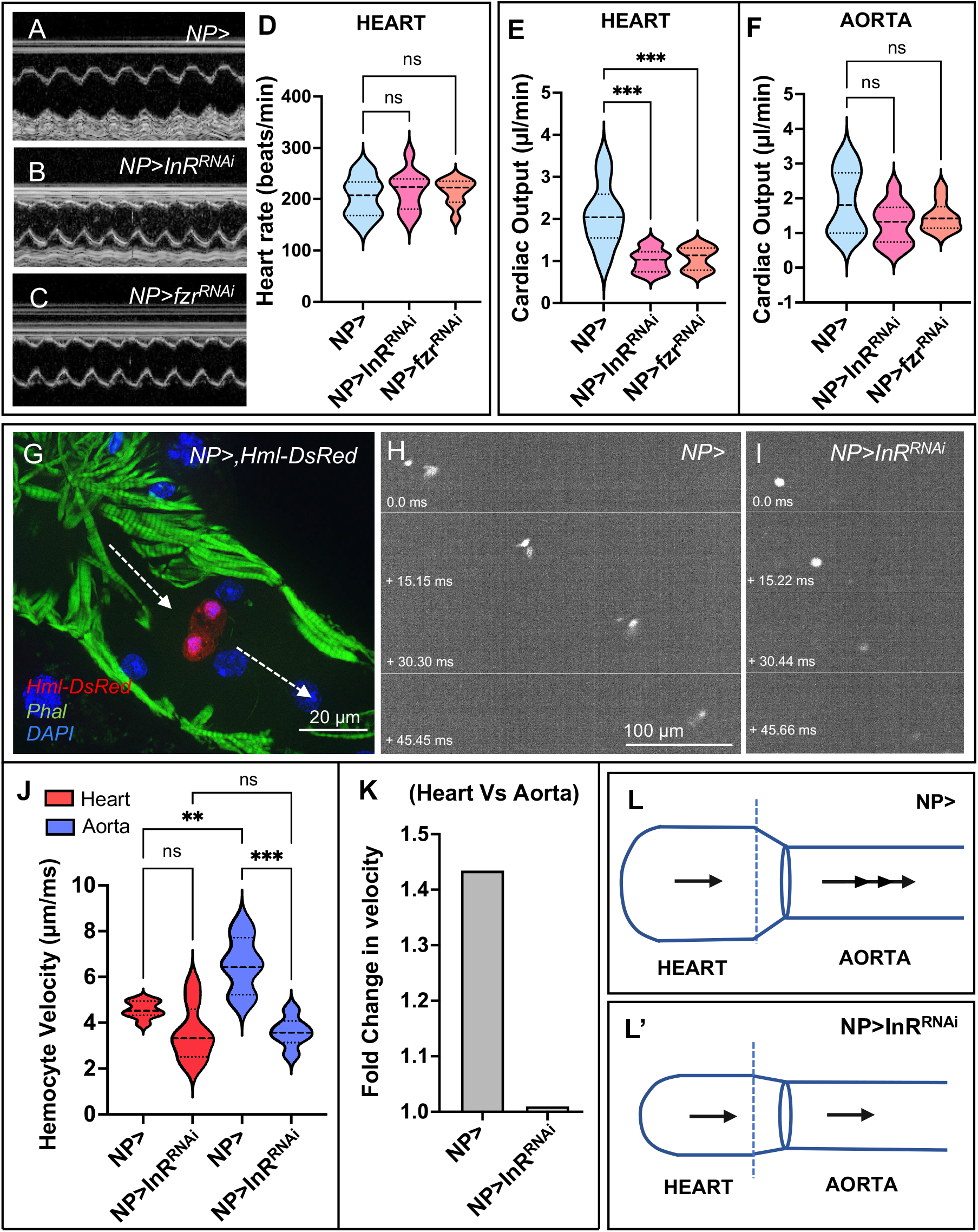
Cardiac output and aorta hemocyte velocity are impacted in cardiac ploidy-reduced animals. **(A-D)** Representative OCT M-mode orthogonal images (see Methods) of heart chamber for control *NP5169-Gal4, UAS-mCherry-NLS* X *w*^*1118*^ (*NP>)* (**A**), *NP>InR RNAi* (**B**) and *NP>fzr RNAi* (**C**). **(D)** Graph of heart rate (beats per min) in the heart chamber of WL3 animals of the indicated genotypes. A two-way ANOVA with multiple comparisons was used to compare heart beats/min for control (*NP>), NP>InR RNAi* and *NP>fzr RNAi* animals and the results are shown as mean ± SD; ^ns^P>0.05. n=10 for each group. Each data set includes two biological repeats. **(E-F)** Graph of cardiac output in the WL3 heart (**E**) and aorta (**F**) chamber of animals of the indicated genotypes. A two-way ANOVA with multiple comparisons was used to compare OCT measurements of heart chamber cardiac output for control (*NP>), NP>InR RNAi* and *NP>fzr RNAi* animals and the results are shown as mean ± SD; ***P <0.001. n= 10 for each group. Each data set includes two biological repeats. **(G)** Fixed image of a control (*NP5169-Gal4, UAS-GFP-NLS* X *w*^*1118*^) WL3 heart chamber (labelled with Phal, green) showing two hemocytes (red, labeled with *Hml-DsRed*). DAPI (blue) labels all nuclei. Arrows point anteriorly, towards the aorta chamber. Scale bar = 100 μm **(H-I)** Representative montage images from movies of WL3 aorta chambers (see Methods) of the indicated genotypes. Anterior is to the right. Hemocytes are labeled with *Hml-DsRed*. Time from the first frame is shown in ms. Scale bar = 100 μm **(J)** Graph showing real time hemocyte velocity in WL3 heart and aorta chambers. A two-way ANOVA with multiple comparisons was used to compare velocity in heart and aorta chambers for control *NP5169-Gal4, UAS-GFP-NLS* X *w*^*1118*^ (*NP>)* and *NP>InR RNAi* animals and the results are shown as mean ± SD; ^ns^P>0.05, **P<0.01, ***P<0.001. n≤5 for each group. Each data set includes two biological repeats. **(K)** Fold change in hemocyte veloicty between heart and aorta chambers is shown for *NP>* and *NP>InR RNAi* animals. **(L-L’)** Schematic illustration depicting chamber dimensions and velocity in control (NP>) and ploidy-reduced (*NP>InR RNAi*) animals. Anterior is to the right.

Given the impact on cardiac output in cardiac ploidy-reduced animals, we next examined the velocity of circulating blood cells (hemocytes) as they move through the cardiac organ (Babcock et al., 2008; Choma et al., 2011). Circulating hemocytes enter the heart chamber at the posterior end and rapidly travel through the aorta with an anterior-directed flow (Babcock et al., 2008) (**Fig5G**). Hemocytes in control animals (*NP>*) accelerate while moving from the heart chamber to the aorta chamber (**Fig5H, J, K, Movie S2**). The average speed of the hemocytes significantly increases from 4.6 µm/ms in the heart to 6.6 µm/ms in the aorta (**Fig5H, J, K**). However, in *NP>InR RNAi* animals, the average velocity of hemocytes remains similar for both chambers: 3.6 µm/ms and 3.5 µm/ms in the heart and aorta, respectively (**Fig5I, J, K, Movie S2**). Overall, our results highlight that hemocytes accelerate as they traverse from the heart to the aorta (**Fig5H, J-L**), but this acceleration does not occur in cardiac ploidy-reduced animals (which have numerous heart chamber defects, **Fig5I, J, K, L’**).

### Chamber-specific ploidy asymmetry is conserved between *Drosophila* and humans

Our findings in *Drosophila* suggest that chamber-specific ploidy regulation in a cardiac organ underlies cardiac organ geometry and function. We next assessed if these findings may be conserved in humans. As we find in *Drosophila*, the human heart becomes polyploid during a mid-point in development (adolescence). The average DNA content of the left ventricular cardiomyocytes increases about 1.7-fold at this time (Bergmann et al., 2015). We measured the nuclear volume of atrial (A) and ventricular (V) cardiomyocytes in adult human samples (see Methods), as an increase in ploidy levels leads to increases in nuclear volume (Cohen et al., 2018; Galitski et al., 1999; Huber and Gerace, 2007; Storchova et al., 2006; Yahya et al., 2022). Tissue sections of explanted donor hearts from 6 subjects (4 females and 3 males, aged in-between 33 to 44, without any history or pathology of heart diseases) were procured from the Duke Human Heart Repository (DHHR, **Fig6A**). Since the left (L) side of the heart (LA and LV) pumps oxygenated blood throughout the human body and the LV chamber wall is the thickest (Ho and Nihoyannopoulos, 2006; Walpot et al., 2019), we compared the nuclear volume of the LV cardiomyocytes to the LA cardiomyocytes. The nuclear volume of LV cardiomyocytes is ∼1.9 fold higher than LA cardiomyocytes (**Fig6B-D**). We did not observe any sex-specific differences in cardiomyocyte nuclear volume (**FigS5A**). These findings suggest that, as in *Drosophila*, the human heart also has chamber-specific asymmetry in nuclear DNA content.

**Figure 6:**
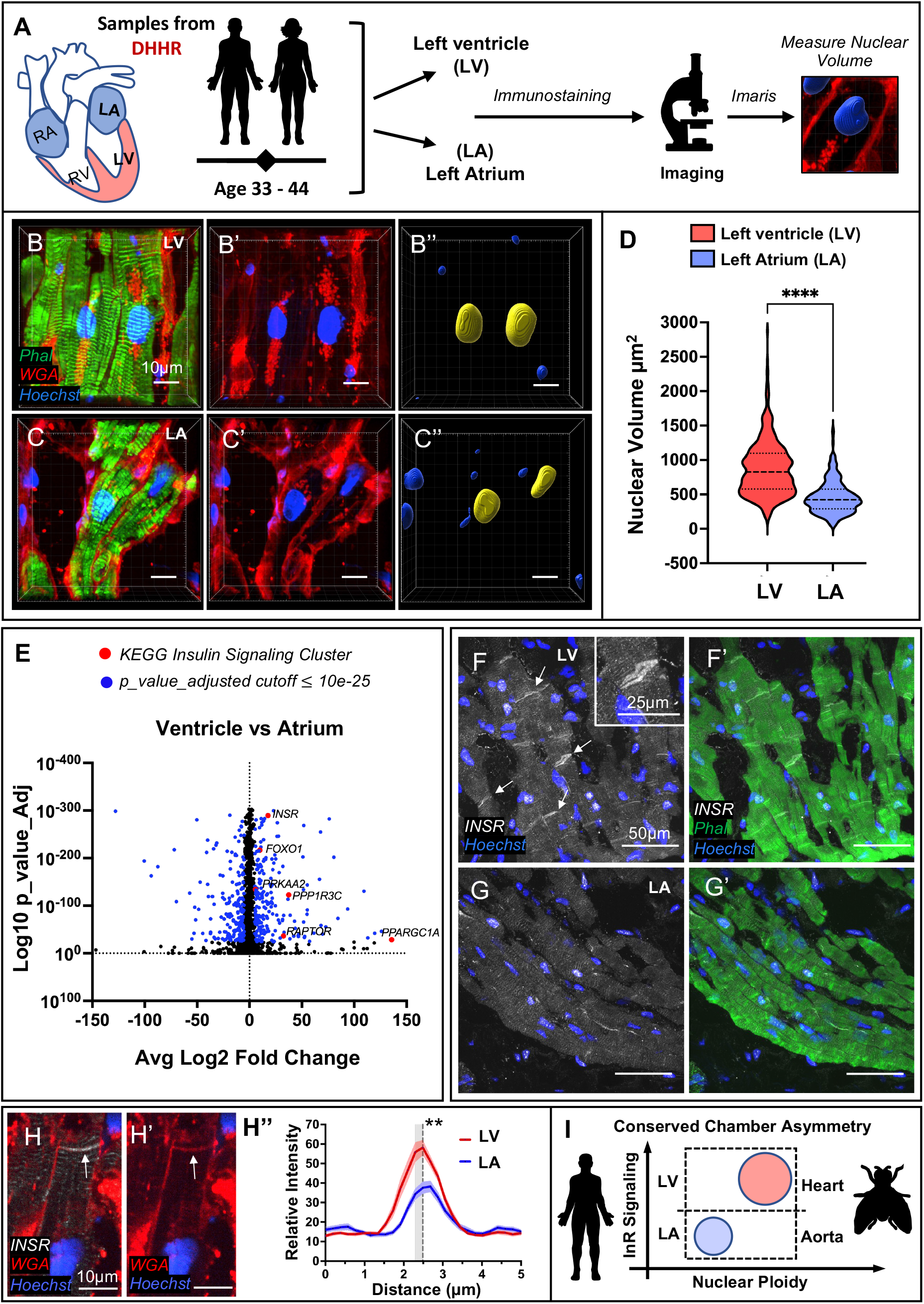
Chamber-specific asymmetry in nuclear volume and insulin signaling in Human hearts. **(A**) Schematic illustration of methods for nuclear volume analysis from human cardiomyocyte tissue sections (see Methods). Tissue sections (10µm thickness) of the left ventricle (LV) and left atrium (LA) from 7 subjects were procured from the Duke Human Heart Repository (DHHR) and nuclear volume was measured using Imaris software. **(B-C’’)** Representative images of LV (**B-B’’**) and LA cardiomyocytes (**C-C’’**) labeled by phalloidin (Phal, green) to show sarcomeres, Wheat Germ Agglutinin (WGA, red) to show cell membranes, and Hoechst (blue) to show nuclei. Nuclear volume of LV cardiomyocytes (**B’’**) and LA cardiomyocytes (**C’’**) were analyzed by Imaris (see Methods). Scale bar = 10 μm. **(D)** Graph of nuclear volume of LV and LA cardiomyocytes. The results are shown as mean ± SD; ****P<0.0001 (Unpaired, two-tailed Student’s t-test). n=6 donor subjects. **(E)** Volcano plot showing differentially expressed genes in ventricular and atrial cardiomyocytes. KEGG analysis (see Methods) show upregulation of insulin signaling pathway as highlighted in red. Dataset retrieved from Litviñuková et.al (2020). **(F-G’)** Representative images of LV (**F-F’**) and LA (**G-G’**) cardiomyocytes labelled with human anti-INSR antibody (white), phalloidin (Phal, green) and Hoechst (blue). Scale bar = 50 μm. Inset in **F** shows a higher magnification view. Inset scale bar= 25 μm. **(H-H’)** Single Z section from an LV cardiomyocyte sample labelled with INSR (white), WGA (red) and Hoechst (blue). Arrows highlight the intercalated regions. Scale bar = 10 μm. **(H’’)** Graph of relative intensity of INSR enrichment at the intercalated regions for LV and LA cardiomyocytes (see Methods). Sidak’s multiple comparisons test was used to compare the fluorescence intensity at the intercalated regions of ventricular and atrial tissue sections. The dashed line indicates the value at which the intensity is significantly different. Result is shown as mean ± SD; **P<0.01. Each data set includes two biological repeats. **(I)** Schematic of the conservation of chamber-specific ploidy asymmetry between *Drosophila* and humans.

Finally, we assessed if there is chamber-specific asymmetry in insulin signaling between ventricular and atrial cardiomyocytes. We first retrieved publicly available combined single cell and nuclear RNA seq datasets (Litvinukova et al., 2020). By pathway analysis (see Methods), we find that insulin signaling is significantly upregulated in the ventricular cardiomyocytes compared to atrial cardiomyocytes (**Fig6E, FigS5B**). Next, to assess these mRNA findings at the protein level, we immunostained LV and LA sections with a human insulin receptor (INSR) antibody. Indeed, LV cardiomyocytes exhibit a distinct INSR enrichment near membrane-associated intercalated discs (Estigoy et al., 2009, **Fig6F, F’, FigS5C-C’**). This pattern is not as prominent in the LA sections (**Fig6G, G’**). By measuring the enrichment of INSR at the intercalated discs, we find that the INSR is significantly higher at this location in the LV cardiomyocytes relative to the LA (**Fig6H-H’’)**. Overall, our results indicate that chamber-specific asymmetry in ploidy is conserved between *Drosophila* and humans and highlight differential insulin signaling as a likely underlying mechanism (**Fig6I)**.

## DISCUSSION

### Developmentally programmed cardiomyocyte ploidy plays a role in heart function and should not be viewed as a regenerative roadblock

Here, we identify temporal and molecular regulation of cardiomyocyte polyploidy in the larval cardiac organ of *Drosophila*. Similar to human cardiomyocytes, which become polyploid during adolescence (Bergmann et al., 2015), polyploidization of *Drosophila* cardiomyocytes occurs during early juvenile (larval) stages. By combining genetics with light microscopy or optical tomography approaches in both living and fixed animals, we highlight multiple defects in cardiac ploidy-reduced animals. Our data strongly argues for an important role of developmentally programmed polyploidy in *Drosophila* larvae. Polyploidy in *Drosophila* cardiomyocytes coincides with larger cell size. Animals with reduced ploidy in the heart chamber exhibit reduced chamber size, stroke volume, cardiac output, and acceleration of circulating hemocytes. Given the high conservation of both transcriptional programming and polyploidy in cardiomyocytes (Akazawa and Komuro, 2005; Olson, 2006; Reim and Frasch, 2010; Stennard and Harvey, 2005), our findings likely apply to any organism with polyploid cardiomyocytes. The idea that programmed polyploidy plays a productive role in cardiac function is not surprising when considering the evolutionary diversity of cardiomyocyte ploidy levels. For example, the giraffe heart is of similar mass as that of other mammals yet generates twice the blood pressure to overcome a severe gravitational challenge. Giraffe left ventricle cardiomyocytes contain, on average, four nuclei and are thus commonly multinucleate polyploid (Ostergaard et al., 2013). Additionally, a large-scale analysis across numerous vertebrate species found that cardiomyocyte polyploidy was likely positively selected for in conjunction with the high metabolic demands of mammalian metabolism (Hirose et al., 2019). Based on our findings here, we argue for further study of the role of developmentally acquired cardiac polyploidy in diverse species.

Blocking developmental polyploidy in mammals is a current strategy aimed at improving heart regeneration (Derks and Bergmann, 2020). Clearly, manipulating developmental cardiomyocyte polyploidy impacts regenerative capacity (González-Rosa et al., 2018; Han et al., 2020; Patterson et al., 2017). However, our findings argue that blocking developmental polyploidy to favor regeneration may ultimately weaken cardiac function. Rather, we argue that developmental polyploidy itself should be disentangled from anti-regenerative gene expression that may arise in conjunction with the transition to the polyploid state. For example, polyploidy and decreased regenerative cell division in mammals is accompanied by higher levels in Thyroid hormone (TH) signaling (Hirose et al., 2019). We note that our KEGG analysis of chamber-specific human heart expression (Litvinukova et al., 2020) also suggests that Thyroid hormone signaling is upregulated in ventricular cardiomyocytes relative to atrial cardiomyocytes (**FigS5B**). In the future, it may be possible to identify TH targets that are specific to either polyploidy or anti-regeneration properties such as inflammatory signaling that produces scarring. Encouragingly, in pigs, overexpression of microRNA-199a successfully uncoupled regeneration and polyploidy. Yet, more work is needed, as this microRNA therapy led to uncontrolled cell proliferation (Gabisonia et al., 2019). Polyploidy is not inherently anti-regenerative, as demonstrated by studies in the mouse liver (Liang et al., 2021; Wilkinson and Duncan, 2021) and many other regenerative organs (Bailey et al., 2021), and roles for polyploid mitosis in developmental organ remodeling in *Drosophila* and mosquito (Fox et al., 2010; Schoenfelder et al., 2014; Stormo and Fox, 2016; Stormo and Fox, 2019). Another important distinction to focus on in the future is between the regulation and function of productive developmentally programmed cardiac ploidy and maladaptive polyploidy after myocardial injury (Beltrami et al., 1997; Gan et al., 2020). We argue that there may be a sweet spot, set by development, of polyploidy level in cardiac organ chambers that promotes optimal heart function.

### Control of ploidy asymmetry as a mechanism to sculpt developing organs

Beyond the polyploid state itself, our data argue that specific levels of polyploidy in specific locations are important in the larval *Drosophila* heart. We show here that differential sensitivity to the Insulin Receptor leads heart chamber cardiomyocytes to undergo speedier endocycles than those of the aorta. As a result, heart cardiomyocytes achieve a higher size and ploidy, and this difference is associated with a wider chamber area in the heart vs. aorta. Notably, we find that hemocytes accelerate when moving from the wider heart to the narrower aorta, in agreement with the Bernoulli principle of fluid dynamics (Badeer, 2001; Bernoulli, 1738). This acceleration is disrupted when heart chamber size is compromised in cardiac ploidy-reduced mutants. We speculate that chamber ploidy asymmetry in fly larvae may function to accelerate and thereby more effectively disperse newly generated clusters of hemocytes throughout the body cavity, to respond to infection more acutely. Future work on the programming of such asymmetry can reveal whether differential insulin signaling is driven by Hox gene patterning differences between the heart and aorta chambers (Bataille et al., 2015; Lo and Frasch, 2001; Lovato et al., 2002; Monier et al., 2005; Perrin et al., 2004; Ponzielli et al., 2002; Schroeder et al., 2022).

In human donor heart samples, we also find asymmetry in chamber ploidy and insulin signaling between the left ventricle (higher ploidy/insulin signaling) and left atrium (lower ploidy/insulin signaling). Our findings suggest conservation of chamber-specific ploidy regulation by insulin signaling. Notably, cardiomyocyte specific ablation of the Insulin Receptor in mice leads to reduced cardiomyocyte size and consequently lower cardiac mass (Belke et al., 2002), and loss of insulin-like growth factor 1 receptor IGR1 in cardiomyocytes leads to disruption of sarcomere structure and intercalated discs (Riehle et al., 2022). The purpose of chamber asymmetry in humans (and mammals in general) may relate to the greater pressure demand on the left ventricle. Our findings here of a 1.8-fold increase in human ventricular vs. atrial nuclear size is consistent with studies in other vertebrates with polyploidy. In quails, hearts ventricular cardiomyocytes are 20% higher in ploidy on average relative to atrial cardiomyocytes (Anatskaya et al., 2001). In mice, where polyploidy is almost exclusively driven by increased nuclear number/myocyte, 77-90% of the ventricular cardiomyocytes are binucleated compared to only 14% in the atrium (Raulf et al., 2015). Connections between insulin and ploidy/cardiomyocyte size may impact understanding of cardiac disease, as patients with type 1 diabetes have reduced ventricular mass without affecting the atrium (Hjortkjær et al., 2019), mirroring our finding of cardiac chamber-specific impacts by the Insulin Receptor in *Drosophila* larvae.

Beyond cardiac biology, our findings here may suggest that ploidy differences along an organ axis may act as a common principle in development to sculpt tissue form and impact tissue function. The unequal growth of body parts is an age-old question in developmental biology (Thompson, 1942). Future work can uncover whether, for example, local ploidy variation accounts for the curvature of polyploid tissues such as the insect egg or for specialized performance of distinct skeletal muscle fibers (Cramer et al., 2020; Windner et al., 2019). Notably, whole genome duplication alters the transcriptome and proteome in non-linear ways (Coate and Doyle, 2010; Maqbool et al., 2010; Yahya et al., 2022; Zhang et al., 2010), highlighting how even a single genome doubling can be transformative in a developmental context. Our work here suggests that future studies should move from a binary (diploid/polyploid) comparisons to take into account the specific level of ploidy. Doing so may accelerate our understanding of the implications of the ubiquitous whole genome duplications in nature.

## METHODS

### Fly Stocks

Flies were raised on standard fly food provided by Archon Scientific. Fly stocks used in this study are: *w*^*1118*^, *hsFLP;tub<FRT>Gal4,UAS-GFP/CyO* (Tub FLP out stock) for clonal analysis, *UAS-mCherry-NLS* (RRID:BDSC_38424, RRID:BDSC_38425), *UAS-GFP-NLS* (RRID:BDSC_4775), *UAS-GFP-NLS*, UAS-Fly-FUCCI (RRID:BDSC_55117, RRID:BDSC_55118) (Zielke et al., 2014b), *UAS-InR RNAi* (VDRC: v992) (Texada et al., 2019), *UAS-fzr RNAi* (VDRC: v25550) (Cohen et al., 2018; Schoenfelder et al., 2014), *UAS-InR*^*CA*^ (RRID:BDSC_8252), *UAS-GFP RNAi* (Gifted by Dr. Zhao Zang) *pnr-Gal4* (RRID:BDSC_3039), *Mef2-Gal4* (RRID:BDSC_27390), *twi-Gal4* (RRID:BDSC_914) and *NP5169-Gal4/cyo* (Kyoto: 113612) (Monier et al., 2005), *TinC-Gal4* (Lo and Frasch, 2001), *HandC-Gal4* (Albrecht et al., 2006), *Hand4*.*2-Gal4 (Han and Olson, 2005)*, and *HandC-GFP* (*Sellin et al., 2006*). *HmlΔ-DsRed* was a gift from Drs. Katja Brükner (Makhijani et al., 2011) and Todd Nystul (UCSF).

Experimental crosses were raised at 25°C, except when heat-shocked at 37 °C for 10-30 min to induce FLP-out clones, or when expressing transgenes, in which case the cross was placed at 29°C. *NP>* is referred to in the text in three contexts: *NP5169-Gal4>UAS-GFP-NLS, NP5169-Gal4>UAS-mCherry-NLS* or *NP5169-Gal4>UAS-mCherry-NLS X w*^*1118*^. Each context is indicated in the corresponding figure legend. For all BrdU feeding experiments, BrdU (100mg/ml, Sigma) was mixed with standard fly food (control food) and fed to larvae of indicated ages for 24 hrs (BrdU Pulse Chase Assay) or for 96-120 hrs (Continuous BrdU Assay). For the restricted diet, embryos were hatched and fed on agar grape juice plates for 72hrs before performing a BrdU pulse chase assay. To measure larval body weight, WL3 larvae were washed in distilled water and measured individually using an analytical balance.

### *Drosophila* Embryo Fixed Imaging

Embryos were fixed and stained by standard protocols (Rothwell and Sullivan, 2007). Briefly, homozygous *UAS-Fly-FUCCI* female flies were crossed to male cardiac mesoderm Gal4 lines (**FigS1A**). Embryos were collected on grape plates with yeast paste for 24hrs at 25°C. After 24hrs, embryos were collected, dechorionated with 50% bleach, and fixed with 4% paraformaldehyde. Fixation buffer was replaced with 1:1 heptane:methanol. Embryos were shaken vigorously, rinsed in methanol, then rinsed in 1X PBS in 0.1% Triton-X (PBST). Embryos were then blocked in 1% goat serum in PBST (block solution) followed by primary antibody incubation in block solution, followed by PBST wash, followed by secondary antibody incubation in block solution. Embryos were mounted in VECTASHIELD (Vector Labs). Primary antibodies used were anti-PH3 (1:1000 mouse) and anti-RFP (1:1000 rabbit). Secondary antibodies used were goat anti-mouse Alexa Fluor 633 (1:500, A-21052; Invitrogen) and goat anti-rabbit Alexa Fluor 568 (1:500, A-11011; Invitrogen). Images were acquired using a Zeiss AxioImager M.2 microscope (20X/0.5 EC Plan-Neofluar objective).

### *Drosophila* Larval Fixed Imaging

Larval heart tissues were dissected in 1X PBS. Due to technical challenges with cleanly recovering the thoracic segments of the dorsal vessel, we used dissection scissors to make an anterior cut at T3. Tissue was fixed and stained by standard protocols (Ponzielli et al., 2002). Briefly, tissue was fixed in 4% paraformaldehyde + 0.3% Triton X-100. Tissues were washed in 1X PBS and blocked in 1X PBS, 0.3% Triton X-100, and 1% normal goat serum. If preforming BrdU immunostaining, tissues were then washed twice in DNAse I buffer (66mM Tris pH 7.5 and 5mM MgCl_2_) for 5 minutes each and then incubated with 10U DNAse (M0303L; New England Biolabs) in 100ul of DNAse buffer at 37 °C for 1hrs. Tissues were then rinsed in 1X PBS in 0.1% Triton-X (PBST) and blocked in 1% goat serum in PBST followed by primary antibody incubation in block solution, followed by PBST wash, followed by secondary antibody incubation in block solution. DAPI (5ug/ul) was added in a final wash step. Laval tissue was mounted in VECTASHIELD (vector labs). For actin labeling, Alexa Fluor 488 & 555 Phalloidin (1:250, 8878, 8953; Cell Signaling) was added together with the secondary antibody in block solution for 2hrs. Primary antibody used was Rat anti-BrdU (1:200, ab6326; Abcam). Secondary antibodies used were goat anti-rat Alexa Fluor 488 (1:500, A-11006; Invitrogen) and goat anti-rat Alexa Fluor 568 (1:500, A-11077; Invitrogen). Images were acquired using an upright Zeiss AxioImager M.2 microscope (20X/0.5 EC Plan-Neofluar objective or 63X/1.4 Oil EC Plan-Neofluar objective).

### Ploidy Analysis of *Drosophila* Cardiomyocytes

Cardiomyocyte ploidy measurements were done as previously described (Clay et al., 2023). Briefly, the larval cardiac organ was dissected in 1X PBS and fixed with 4% formaldehyde solution. Dissected hearts were then washed with 1X PBS and then transferred to a siliconized cover slip. A charged slide was gently placed on top of the siliconized cover slip with a drop (∼10ul) of 1X PBS. Pressure was gently applied to all corners of the cover slip and then further pressure was applied using a vise. The glass slide was submerged into liquid nitrogen and then using a razor blade the siliconized cover slip was quickly removed. The slides were then transferred to a Coplin jar containing -20°C 90% EtOH. Samples were air dried and rinsed with 1X PBS before labeling with DAPI (5µg/ml) for 10 minutes. Samples were then rinsed twice with 1X PBS and then mounted in VECTASHIELD (vector labs). For ploidy measurement, testes from adult flies were dissected and processed together with the dissected hearts on the same slide. Z stack images of 1µm thickness were acquired using an upright Zeiss AxioImager M.2 (63X/1.4 Oil EC Plan-Neofluar objective or 40X/ 1.3 Oil EC Plan-Neofluar objective). Ploidy analysis was done using Fiji. Ploidy of each nucleus was calculated to the median DAPI intensity of the haploid spermatids imaged on the same slide with the same setting. All the cardiomyocyte nuclei were labeled with mCherry-NLS (*NP5169-Gal4>UAS-mCherry-NLS*) to distinguish them from other the nonspecific nuclei (such as hemocytes, plasmocytes, pericardial cells). Note that the four cardiomyocytes of the apex (A7 segment) were excluded from our heart chamber ploidy plots (**Fig 2A, 3G**) because of their distinct cell shape.

### OCT Measurements of *Drosophila* Larval Hearts

Cardiac function of *Drosophila* WL3 cardiac organs was measured using Thorlabs’ Ganymede™ OCT Systems as previously described (Wolf et al., 2006; Yu et al., 2013; Yu et al., 2015). To image live larval hearts, animals were quickly rinsed in distilled water and transferred to an adhesive platform made from transparent tape (3M Heavy Duty Scotch tape) attached to a glass slide. Each larva was placed ventral side down, with the dorsal side exposed for OCT imaging. Videos of the posterior heart and anterior aorta were acquired using the ThorImage OCT 4.2. Each video was acquired at a speed of 100 frames/sec for 5 seconds duration. Multiple M-Mode transverse and sagittal real-time videos were recorded for the heart and the aorta chamber. As the larval cardiac organ lumen was not exactly circular, we directly calculated the area (a) of the lumen using Fiji software. To do so, End Systolic Dimension (ESD= a_S_) and End Diastolic Dimension (EDD= a_D_) were calculated by tracing the lumen area from the transverse OCT images. Each chamber length (*l*) was measured from the sagittal M-Mode OCT images directly using the ThorImage OCT 4.2 software. For measuring the total chamber wall thickness during diastole, anterior (*t*_*A*_) and posterior (*t*_*P*_) wall thicknesses were measured using Fiji Software and summed. The number of heart beats for each chamber was recored from the transverse M-mode OCT videos using orthogonal view of stacks in Fiji. Heart beats per second were calculated by dividing the number of heart beats with the length of the video (i.e. 5 seconds). Heart rate is defined as number of heart beats per minute. To calculate the heart rate, the number of heart beats per second was multiplied by 60 (**Fig5D, FigS4N**).

The stroke volume and cardiac output of each chamber was measured as previously described (Klassen et al., 2017, **FigS2I**). To calculate the stroke volume of each chamber, the EDD area was subtracted from the ESD area and then multiplied by the chamber length. The cardiac output of each chamber was measured by multiplying the chamber heart rate by the chamber stroke volume.

### Live Imaging of *Drosophila* Larval Hearts

To visualize the chamber specific movement of the hemocytes inside the larval cardiac organ, we imaged the fluorescently labeled hemocytes (*Hml*Δ*-DsRed)* (Makhijani et al., 2011) in live animals using an Andor Dragonfly 505 unit with a Borealis illumination spinning disk confocal and a Zyla PLUS 4.2 Megapixel sCMOS camera and 20x/0.75 HC PL APO CS2 (Leica 11506517), Dry Objective, WD: 0.62 mm. To restrict the moment of larvae, we used the adhesive platform described above. Each larva was placed ventral side down, with the dorsal side exposed for imaging using the Andor platform. Furthermore, to identify the cardiac organ while imaging, it was fluorescently labeled (*NP5169>UAS-GFP-*NLS). Multiple videos were recorded for each animal to determine the chamber specific movement of the hemocytes.

### Human Tissue Sample Preparation

Flash frozen left ventricular (LV) and left atrial (LA) tissue samples from explanted hearts of 6 subjects were procured from the Duke Human Heart Repository (DHHR) under the review of Duke University Health System (DUHS) institutional review board (IRB) (Pro00005621). All subjects were as follows: (Male 1: age 42, Male 2: age 33, Male 3: age 41, Female 1: age 42, Female 2: age 44, and Female 4: age 44. Tissue samples were mounted in O.C.T. compound (Scigen #4586) and sectioned at 10µm thickness with the help of the Duke Substrate Service Core & Research Support (SSCRS). Each section was transferred to a positively charged glass slide, labeled and stored at -80°C before immunostaining.

### Human Tissue Immunostaining

Frozen tissue sections were thawed at room temperature for 10 minutes (min) and then washed twice with 1X PBS for 5 min each to dissolve the O.C.T. medium. Tissue sections were then fixed with 4% paraformaldehyde solution for 15 min at room temperature (RT). Samples were then washed twice with 1X PBS and incubated with Alexa Fluor 633 conjugate of Wheat germ agglutinin (WGA) (1:250, W21404; Invitrogen) in 1X PBS for overnight at 4°C. Note that WGA staining was performed before permeabilization using Triton X-100. To wash off the WGA solution, samples were then washed thrice with PBST (1X PBS containing 0.1% Triton X-100) for 5 min each and then incubated in blocking solution (5% Serum in PBST) for 40 minutes at RT. Samples were then incubated with primary antibody in blocking solution overnight at 4°C. Tissues samples were then washed thrice with PBST for 10 mins each and then incubated with secondary antibody, Hoechst and Alexa Fluor 488 Phalloidin (1:250, 8878; Cell Signaling) in block solution at RT for 2hrs. Tissues were then washed thrice with PBST for 10 mins each and mounted in VECTASHIELD (vector labs). For nuclear volume analysis, tissue sections were labeled first with WGA in 1X PBS, overnight and then incubated with Phalloidin and Hoechst in 1X PBS for two hours. The primary antibody used for this study was rabbit anti-Insulin Receptor (1:200, ab137747; Abcam). The secondary antibody used was goat anti-rabbit Alexa Fluor 546 (1:500, A-11006; Invitrogen).

### Human Tissue Image Acquisition and Nuclear Volume Analysis

For nuclear volume analysis, Z-stack images of each tissue section were imaged using an Andor Dragonfly 505 unit with a Borealis illumination spinning disk confocal, with a Z-step size of 0.5µm. Samples were imaged using an Andor Zyla PLUS 4.2 Megapixel sCMOS camera together with a 63x/1.47 TIRF HC PL APO CORR (Leica 11506319), Oil objective, WD: 0.10 mm. For analysis of nuclear volume of cardiomyocytes, we used the Imaris Version 8.2 module ImarisCell. The protocol for analyzing the cardiomyocyte nuclear volume was adapted from (Bensley et al., 2016). Briefly, using ImarisCell, the nuclear volume of all nuclei in the Z stack were recorded. Using phalloidin to identify sarcomere-containing cells and WGA labeling to identify cell borders, cardiomyocyte nuclei were selected manually. Only mononucleated cells were imaged, and binucleated cells were excluded. About 50 to 100 nuclei were processed for each group from multiple images.

Fiji software was used to analyze the fluorescence intensity of INSR signal. Using the Plot Profile program in the Analysis Section, we measured the intensity of INSR at the intercalated regions of a single Z-section that had the highest enrichment of signal. Intensity profiles of 30 cells from multiple images were calculated and data were plotted as previously described (Sawyer et al., 2017).

### Single-cell Data RNA-seq Analysis

“Heart Global” data sets (accession no: ERP123I38) containing single cell and single nuclei RNA-Seq data were downloaded from the Heart Cell Atlas (https://www.heartcellatlas.org) (Litvinukova et al., 2020). Differential gene expression between ventricular cardiomyocytes (LV+RV) and atrial cardiomyocytes (LA+RA) was done using the Seurat R package (Hao et al., 2021). The list of differentially expressed genes between ventricular and atrial cardiomyocytes are summarized in **Supplemental Dataset 1**, in the sheet named “DE_V_vs_A”. Upregulated genes in ventricular cardiomyocytes are summarized in **Supplemental Dataset 1**, in the sheet named “V_UP”. Pathway analysis was done using DAVID (Huang da et al., 2009; Sherman et al., 2022) online software. A list of KEGG upregulated pathways in ventricular cardiomyocytes is summarized in **Supplemental Dataset 1**, in the sheet named “KEGG _V_UP”.

### Quantification and Statistical Analysis

Statistical methods of analysis, number of biological replicates, values of **n**, and P values are detailed in the figure legends. Statistical analysis was performed using GraphPad Prism 9.3.1. Statistical notations used in figures: (P > 0.05, (ns) not significant); (P < 0.05, *); (P < 0.01, **); (P < 0.001, ***); (P ≤ 0.0001, ****).

## ACKNOWLEDGMENTS

We thank the Bloomington Drosophila Stock Center and Vienna Drosophila Resource Center for providing the reagents used in this study. We thank Ruth Montague, Dr. Jessica Sawyer, Dr. Matt Andrusiak, and Rebeccah Stewart for assistance. We thank Dr. Zhao Zhang and Dr. Sarah Goetz for providing fly and chemical reagents. We thank Dr. Lisa Cameron, Dr. Yashen Gao and the Duke Light Microscopy Core for image acquisition assistance. We thank the Duke Human Heart Repository (DHHR) for proving the human heart donor tissue samples. We thank Paula Newell, Cameron Leonard and Substrate Service Core & Research Support (SSCRS) for providing assistance with sectioning of the human tissue samples. We thank Drs. Sarah Goetz and Zhao Zhang and Fox lab members for providing comments on the manuscript.

This project is supported by an American Heart Association (AHA Award ID 23POST1013432) and a Duke Regeneration Center (DRC)-Postdoctoral Research Award to Accelerate Career Independence to A. Chakraborty, an NIH predoctoral fellowship (F31HL162460) to S. DeLuca., a National Heart, Lung, and Blood Institute grant (R01HL164013) to N. Bursac, a National Heart, Lung, and Blood Institute grant (R01HL158718) to M. Wolf, and a National Institute of General Medical Sciences grant (R01GM118447) to D.T. Fox. Prior early-stage support was also provided by an AHA Scientist Development grant to D.T. Fox (15SDG24480160) and a National Heart, Lung, and Blood Institute grant to N.G. Peterson (F31HL140811).

## SUPPLEMENTAL MATERIAL

### Supplementary Figure Legends

**Figure S1:**
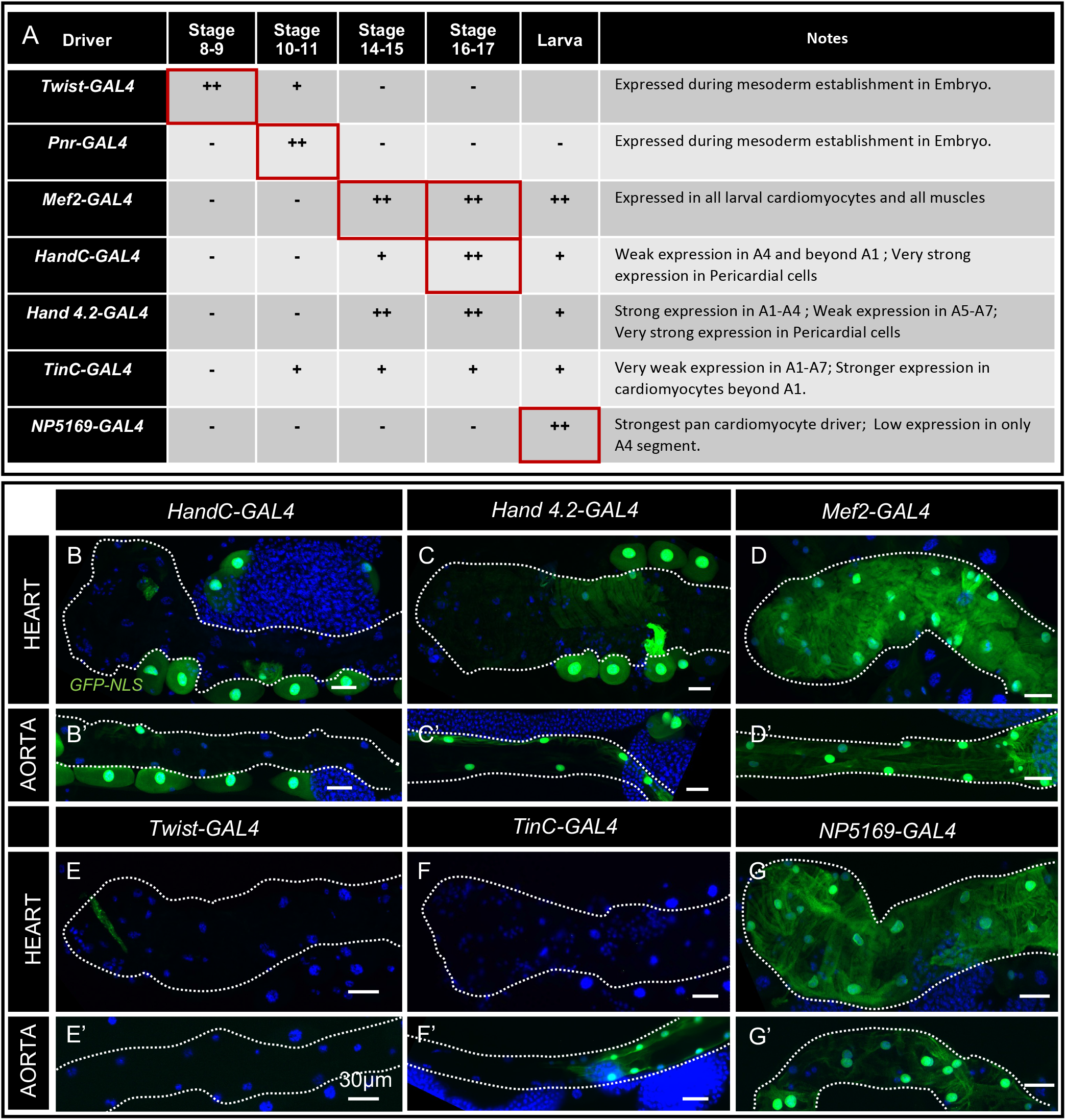
Expression of cardiac Gal4 drivers in the embryo and larva. **(A)** Summarized expression pattern of selected Gal4 drivers in embryos and larvae. Red boxes indicate the Gal4 driver used for stage-specific expression in embryos and larvae. **(B-G’)** Representative images showing expression patterns (UAS-GFP, green) of *HandC-GAL4* (**B-B’**), *Hand4*.*2 GAL4* (**C-C’**), *Mef2-GAL4* (**D-D’**), *twi-GAL4* (**E-E’**), *TinC*-GAL4 (**F-F’**) and *NP5169-Gal4* (**G-G’**) in the WL3 heart and aorta. Nuclei in all images are labeled with DAPI (blue). Scale bar = 30 μm.

**Figure S2:**
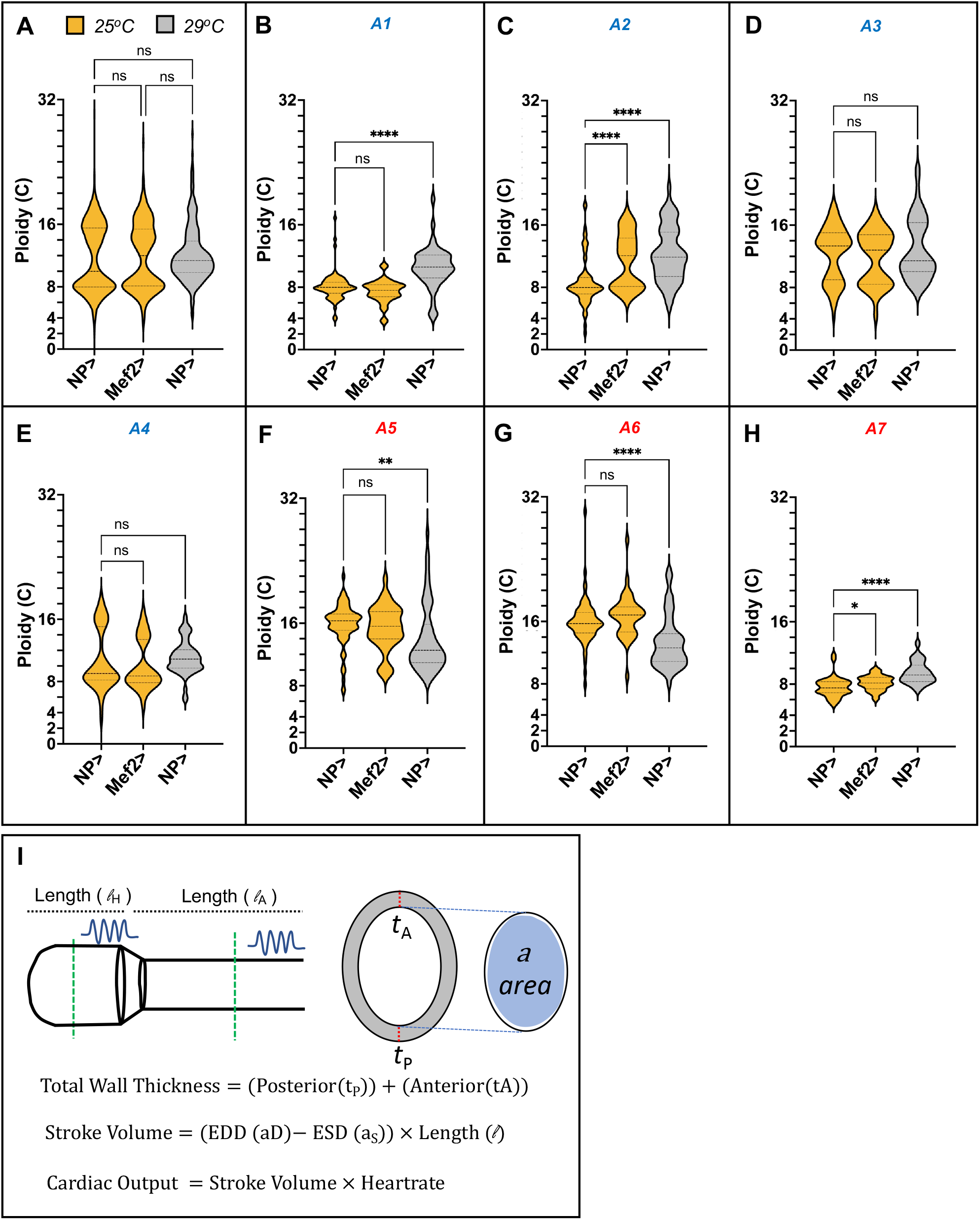
Segmental and temperature regulation of cardiomyocyte ploidy. **(A)** Graph showing temperature-specific distribution of total WL3 cardiomyocyte ploidy throughout the dorsal vessel (segments A1-7). A two-way ANOVA with multiple comparisons was used to compare total ploidy among *NP5169-Gal4>UAS-mCherry* (*NP>)* at 25°C, *Mef2-Gal4>UAS-mCherry* (*Mef2>)* at 25°C and *NP5169-Gal4>UAS-mCherry* (*NP>)* at 29°C and the results are shown as mean ± SD; ^ns^P>0.05. n= 10 for each group. **(B-H)** Graphs showing temperature-specific distribution of WL3 cardiomyocyte ploidy in segments A1 to A7. A two-way ANOVA with multiple comparisons was used to compare total ploidy among *NP>* at 25°C, *Mef2>* at 25°C and *NP>* at 29°C and the results are shown as mean ± SD; ^ns^P>0.05, *P<0.05, **P<0.01, ****P<0.0001. n= 10 for each group. Each data set includes two or more biological repeats. **(I)** Schematic illustration of the parameters measured in the cardiac organ using OCT analysis.

**Figure S3:**
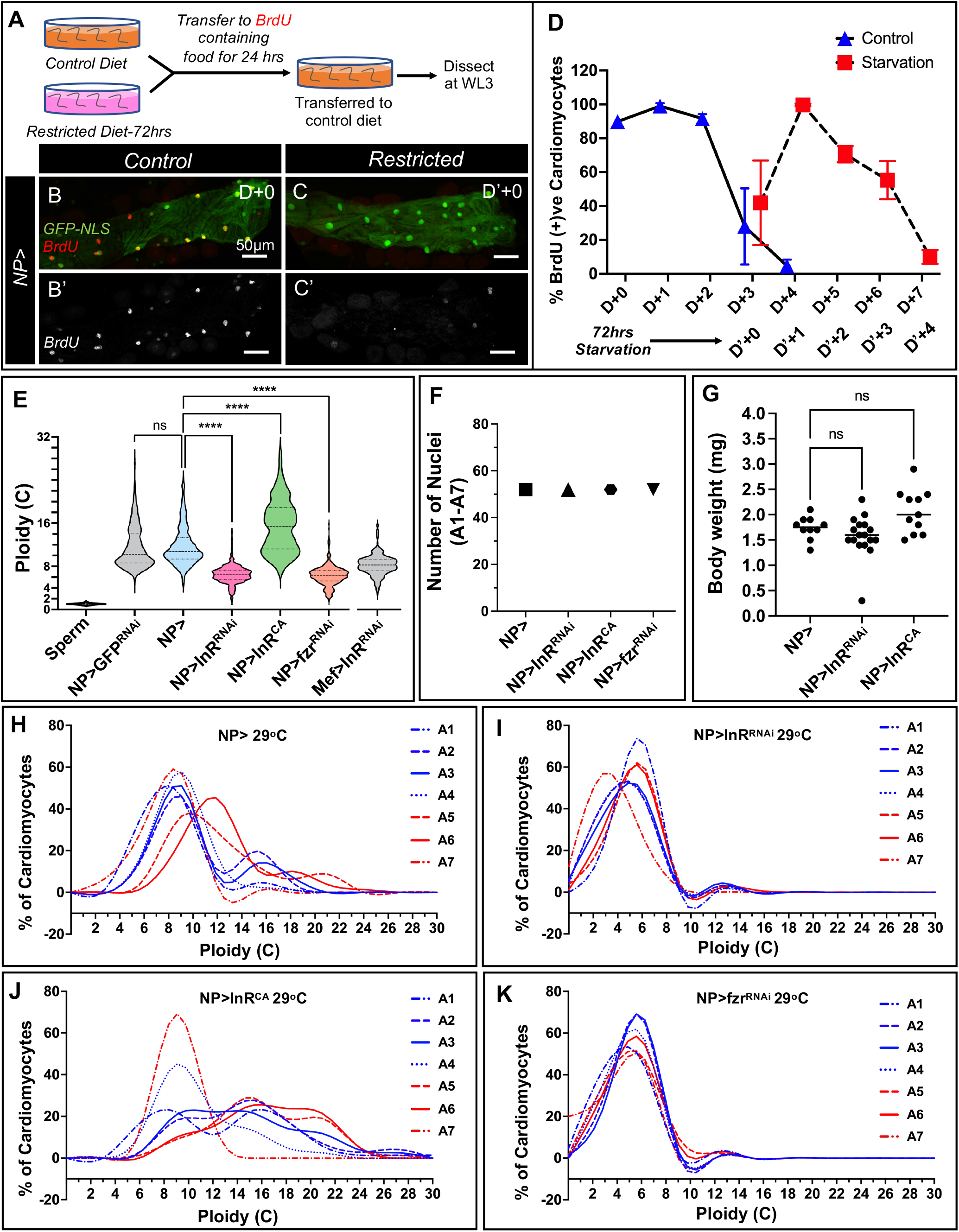
Data supporting the conclusion that larval cardiomyocyte endocycles are regulated by insulin signaling. **(A)** Schematic representation of BrdU pulse chase assay performed on fully fed animals (control diet) vs. animals on a restricted diet for 72hrs (Methods). (**B-C’**) Representative images of BrdU positive (red) cardiomyocytes (green) in the heart chamber at WL3 after being fed BrdU during D+0 (first 24 hours after hatching) (**B-B’**) and after 72hrs of restricted diet at D’+0 (**C-C’**). *NP5169-Gal4>UAS-GFP-NLS* labels cardiomyocytes. Scale bar = 50 μm. **(D)** Graph showing the percentage of BrdU positive WL3 cardiomyocytes from animals fed BrdU for a 24-hour period. Animals were either continuously reared on a control diet or were first on a restricted diet for 72 hours as shown in (**A**). The X-axis indicates the 24-hour period during which the animals were fed BrdU. n=5 for each group. Each data set includes two biological repeats. All crosses were maintained at 25°C. **(E)** Graph showing total WL3 cardiomyocyte ploidy (A1-A7) in animals of the indicated genotypes. A two-way ANOVA with multiple comparisons was used to compare the total ploidy among control *NP5169-Gal4, UAS-mCherry-NLS* X *w*^*1118*^ (*NP>), NP>GFP RNAi, NP>InR RNAi, NP>InR*^*CA*^ and *NP>fzr RNAi* animals and the results are shown as mean ± SD; ****P<0.0001. n= 10 for each group. Each data set includes two or more biological repeats. **(F)** Graph showing the total number of WL3 cardiomyocyte nuclei in segments A1 to A7 (52 cardiomyocytes) in animals of the indicated genotypes. n= 10 for each group. Each data set includes two biological repeats. **(G)** Graph showing WL3 body weight (see Methods) of animals of the indicated genotypes. A two-way ANOVA with multiple comparisons was used to compare the total ploidy among control *NP5169-Gal4, UAS-mCherry-NLS* X *w*^*1118*^ (*NP>), NP>InR RNAi* and *NP>InR*^*CA*^ animals and the results are shown as mean ± SD; ^ns^P>0.05. n= 10 for each group. Each data set includes two biological repeats. **(H-K)** Histograms showing WL3 cardiomyocyte ploidy distribution in each segment (A1-A7) for animals of the indicated genotypes, raised at 29 °C. n=10 for each group. Each data set includes two or more biological repeats.

**Figure S4:**
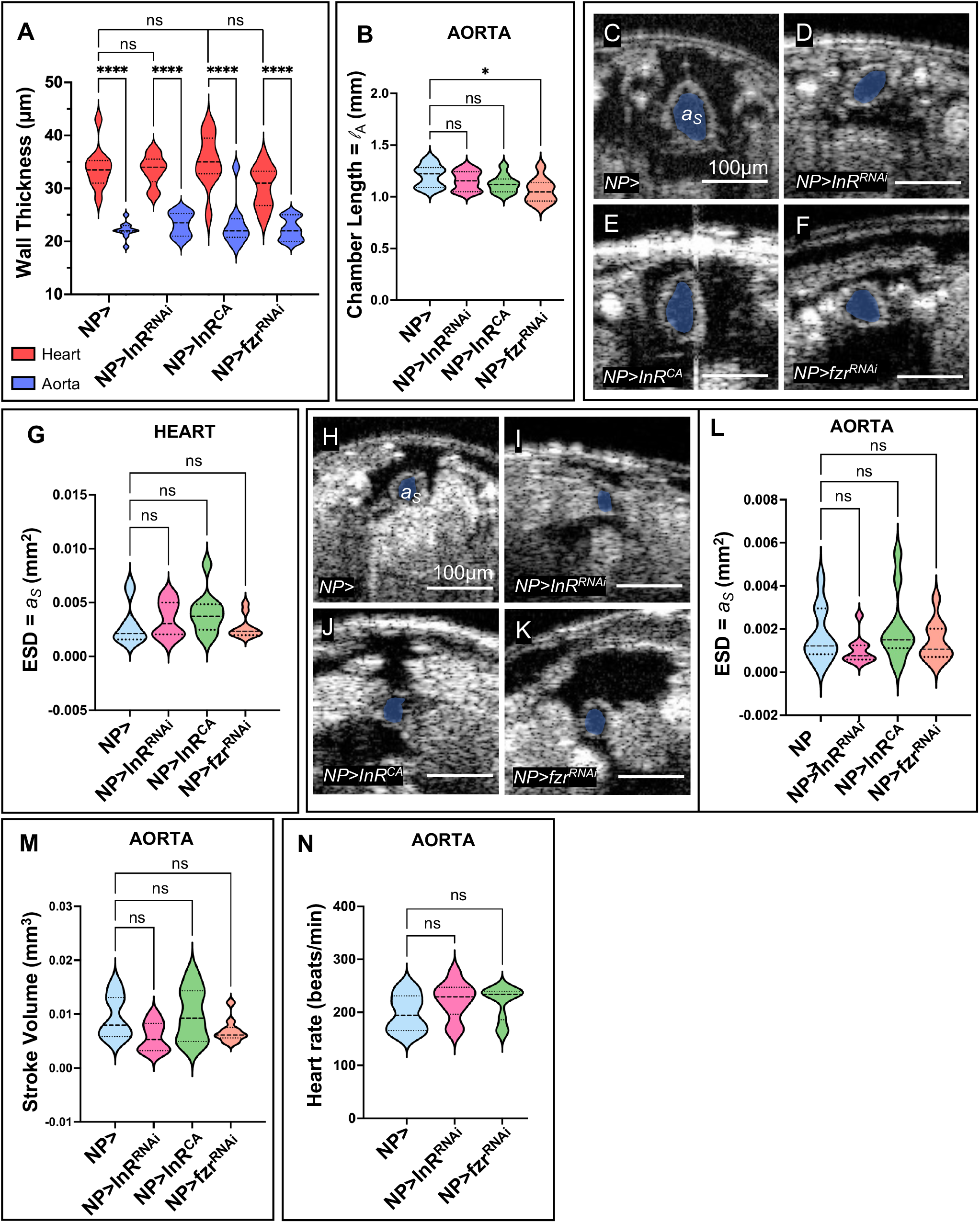
Aorta chamber dimensions are less affected in mutants with altered ploidy. **(A)** Graph of heart wall thickness in the heart and aorta chambers of WL3 animals of the indicated genotypes. A two-way ANOVA with multiple comparisons was used to compare OCT measurements of heart aorta chamber wall thickness for control *NP5169-Gal4, UAS-mCherry-NLS* X *w*^*1118*^ (*NP>), NP>InR RNAi, NP>InR*^*CA*^ and *NP>fzr RNAi* animals and the results are shown as mean ± SD; ^ns^P>0.05, ****P<0.0001. n= 10 for each group. Each data set includes two biological repeats. **(B)** Graph of WL3 aorta chamber length in animals of the indicated genotypes. A two-way ANOVA with multiple comparisons was used to compare OCT measurements of aorta chamber length for control *NP5169-Gal4, UAS-mCherry-NLS* X *w*^*1118*^ (*NP>), NP>InR RNAi, NP>InR*^*CA*^ and *NP>fzr RNAi* animals and the results are shown as mean ± SD; *P<0.05. n= 10 for each group. Each data set includes two biological repeats. **(C-F)** Representative transverse two-dimensional real-time OCT images of WL3 heart chamber End Systolic Dimension are (ESD, a_S_, pseudo colored in gray) for control *NP5169-Gal4, UAS-mCherry-NLS* X *w*^*1118*^ (*NP>)* (**C**), *NP>InR RNAi* (**D**), *NP>InR*^*CA*^ (**E**) and *NP>fzr RNAi* (**F**). Scale bar = 100 μm. **(G)** Graph showing WL3 heart chamber a_S_ for animals of the indicated genotypes. A two-way ANOVA with multiple comparisons was used to compare heart chamber a_S_ area among control (*NP>), NP>InR RNAi, NP>InR*^*CA*^ and *NP>fzr RNAi* animals and the results are shown as mean ± SD; ^ns^P>0.05. n= 10 for each group. Each data set includes two biological repeats. **(H-K)** Representative transverse two-dimensional real-time OCT images of WL3 aorta chamber a_S_ pseudo-colored in gray for control *NP5169-Gal4, UAS-mCherry-NLS* X *w*^*1118*^ (*NP>)* (**H**), *NP>InR RNAi* (**I**), *NP>InR*^*CA*^ (**J**) and *NP>fzr RNAi* (**K**). Scale bar = 100 μm. **(L)** Graph of WL3 aorta a_S_ for animals of the indicated genotypes. A two-way ANOVA with multiple comparisons was used to compare OCT measurements of WL3 aorta chamber a_S_ for control (*NP>), NP>InR RNAi, NP>InR*^*CA*^ and *NP>fzr RNAi* animals and the results are shown as mean ± SD; ^ns^P>0.05. n= 10 for each group. Each data set includes two biological repeats. **(M)** Graph of WL3 stroke volume of the aorta chamber for animals of the indicated genotypes. A two-way ANOVA with multiple comparisons was used to compare OCT measurements of aorta chamber stroke volume for control (*NP>), NP>InR RNAi, NP>InR*^*CA*^ and *NP>fzr RNAi* animals and the results are shown as mean ± SD; ^ns^P>0.05. n= 10 for each group. Each data set includes two biological repeats. **(N)** Graph of heart rate (beats per min) in the aorta chamber of WL3 animals of the indicated genotypes. A two-way ANOVA with multiple comparisons was used to compare heart beats/min for control *NP5169-Gal4, UAS-mCherry-NLS* X *w*^*1118*^ (*NP>), NP>InR RNAi* and *NP>fzr RNAi* animals and the results are shown as mean ± SD; ^ns^P>0.05. n=10 for each group. Each data set includes two biological repeats.

**Figure S5:**
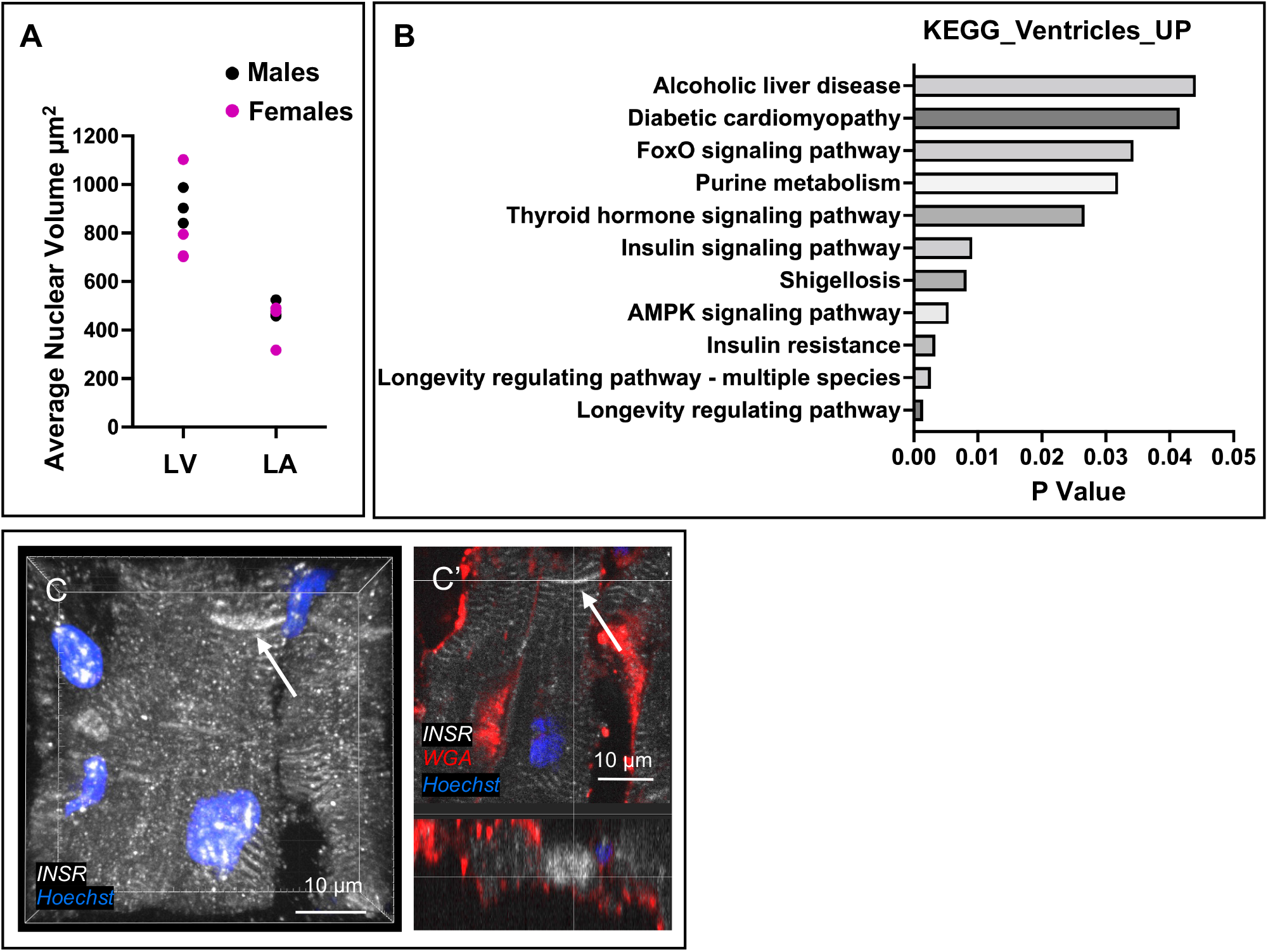
Supporting evidence of asymmetry in nuclear volume and insulin signaling between the Human left ventricle and atrium. **(A)** Graph of nuclear volume of LV and LA cardiomyocytes for males and females. Males were of ages 42 years, 33 years and 41 years. Females were of age 42 years, 44 years, 42 years and 44 years. **(B)** KEGG analysis of the upregulated genes in ventricles from Litviñuková et.al (2020). p value<0.05. **(C-C’)** Z stack image of LV cardiomyocyte (**C**). Arrowhead highlighting INSR enrichment pattern (**C**). Arrowhead highlighting INSR enrichment at the membrane near intercalate region (**C’**). Scale bar = 10 μm.

**Supplemental Dataset 1: Differentially expressed genes in Ventricular vs Atrial cardiomyocytes**. Sheet DE_V_vs_A lists differentially expressed genes in ventricular and atrial cardiomyocytes. Sheet V_UP lists genes upregulated in ventricular cardiomyocytes. Sheet KEGG_V_UP lists gene pathways upregulated in ventricular cardiomyocytes. Dataset retrieved from Litviñuková et.al (2020). **Related to Figure 6E**.

**Supplemental Movie 1:** Ploidy **reduced mutants show a smaller lumen area**.

M-Mode sagittal real-time videos of the heart chamber in control NP> (*NP5169-Gal4, UAS-GFP-NLS* X *w*^*1118*^), *NP>InR-RNAi, NP>InR*^*CA*^ and *NP>fzr-RNAi* animals. Related to **Figure 4A-E**.

**Supplemental Movie 2: Hemocyte velocity in the aorta decreases in *InR* knockdown animals**. Movement of hemocytes in the aorta for control *NP> (NP5169-Gal4, UAS-GFP-NLS* X *w*^*1118*^) and *NP>InR-RNAi* animals. Hemocytes are labeled with *Hml-DsRed*. Related to **Figure 5H-K**.

